# Shared B cell memory to coronaviruses and other pathogens varies in human age groups and tissues

**DOI:** 10.1101/2020.12.01.407015

**Authors:** Fan Yang, Sandra C. A. Nielsen, Ramona A. Hoh, Ji-Yeun Lee, Tho D. Pham, Katherine J. L. Jackson, Krishna M. Roskin, Yi Liu, Robert S. Ohgami, Eleanor M. Osborne, Claus U. Niemann, Julie Parsonnet, Scott D. Boyd

## Abstract

Vaccination and infection promote the formation, tissue distribution, and clonal evolution of B cells encoding humoral immune memory. We evaluated convergent antigen-specific antibody genes of similar sequences shared between individuals in pediatric and adult blood, and deceased organ donor tissues. B cell memory varied for different pathogens. Polysaccharide antigen-specific clones were not exclusive to the spleen. Adults’ convergent clones often express mutated IgM or IgD in blood and are class-switched in lymphoid tissues; in contrast, children have abundant class-switched convergent clones in blood. Consistent with serological reports, pre-pandemic children had class-switched convergent clones to SARS-CoV-2, enriched in cross-reactive clones for seasonal coronaviruses, while adults showed few such clones in blood or lymphoid tissues. These results extend age-related and anatomical mapping of human humoral pathogen-specific immunity.

**One Sentence Summary:** Children have elevated frequencies of pathogen-specific class-switched memory B cells, including SARS-CoV-2-binding clones.

## Main Text

Clonal proliferation of lymphocytes is critical for the adaptive immune system to respond to pathogens. The clonal identity of a B cell can be traced by the sequence of its B cell receptor (BCR), which determines its antigen specificity (*1–3*). Immunoglobulin (Ig) sequences are formed via irreversible V(D)J gene segment rearrangement and can be diversified through somatic hypermutation (SHM) (*4*) and class-switch recombination (CSR) (*5*). Convergent or “public” Ig sequences showing similar antibody gene rearrangements can be found in the primary BCR repertoires of naive B cells, owing to biases in V(D)J gene recombination mechanisms, but are enriched in individuals exposed to the same antigen, reflecting an antigen-driven selection for particular sequence features in the Ig heavy (IGH) or Ig light chain complementarity-determining region (CDR) loops of the BCR (*6–12*). These Ig sequences, when detected in antigen-experienced B cells, document previous infections and allow for the reconstruction of the immunological memory of an individual. Recently, high-throughput sequencing (HTS) has been applied to study convergent IGH repertoires across individuals in response to vaccination and infection (*11–14*). It is still unclear, however, how our immune memories to different antigens distribute in immunological tissues and change during an individual’s lifespan.

The immune system gradually matures and accumulates immunological memory from infancy onward (*15*). The immune responses to some pathogens appear to differ between young children and adults. For example, young children infected with human immunodeficiency virus (HIV) can achieve neutralizing antibody breadth relatively early after infection, via different mechanisms than adults (*16*). Data emerging from the current coronavirus disease 2019 (COVID-19) pandemic show that children usually have milder symptoms than adults (*17–22*), potentially due to differences of viral receptor expression and other host factors, but also, as a recent serological study suggests, elevated pre-pandemic frequencies of *severe acute respiratory syndrome coronavirus 2* (SARS-CoV-2) binding and neutralizing antibodies in children, stimulated by other coronavirus infections (*23, 24*). During the COVID-19 pandemic, infected children compared to adults have lower SARS-CoV-2 antibody titers and a predominance of IgG antibodies specific for the S protein but not the N protein, with these differences attributed to faster clearance of virus limiting the amount of viral antigens produced in the body (*25*). The ways in which B cell clonal populations specific for coronaviruses and other pathogens may differ between children and adults is unknown. Most studies of human B cells rely on peripheral blood mononuclear cells (PBMCs), which represent a small part of an individual’s full IGH repertoire, with secondary lymphoid tissues in lymph nodes, spleen, and the gastrointestinal tract harboring larger numbers of B cells and being major sites for SHM and CSR (*26, 27*). Specialized immune responses in some lymphoid tissues have been reported; for example, antibody responses to polysaccharide antigens are decreased in the absence of functional splenic tissue (*28–33*). It is still unclear if B cell clones are widely dispersed in different secondary lymphoid tissues or enriched in particular tissues depending on their antigen or pathogen specificity.

To better understand how the antigen-specific B cell memory compartment changes through the human lifespan, and distributes itself across immunological tissues, we systematically characterized the convergent IGH repertoires specific to six common pathogens or vaccine types, and two viruses not encountered by the participants (Ebola virus and SARS-CoV-2) in pre-COVID-19 pandemic individuals from newborns to 87 years of age, and in tissues including cord blood, peripheral blood, spleen, and lymph nodes. For blood B cell analysis, we analyzed 12 cord blood samples, 93 peripheral blood samples from 51 children (*34*), 122 healthy human adult peripheral blood samples (*34*), and eight blood samples from deceased organ donors (table S1)). All children were known to be exposed to *Haemophilus influenzae* type b (Hib), *Pneumococcus pneumoniae* (PP), tetanus toxoid (TT), and influenza virus (Flu) antigens via vaccination, and it is highly likely that all were exposed to respiratory syncytial virus (RSV) in their first three years of life (*35*). The children did not receive *Neisseria meningitidis* (NM) vaccination. The detailed vaccination histories of the adults are unknown. To identify convergent IGH sequences for the six pathogens in blood samples, we clustered IGH reads with known pathogen-specific reference IGH sequences (table S2) based on their IGHV and IGHJ gene segment usage, CDR-H3 length, and at least 85% CDR-H3 amino acid sequence similarity (see Methods).

We hypothesized that cord blood samples would show the least evidence of convergent pathogen-specific IGH gene sequences, if these IGH were in fact truly stimulated by pathogen or vaccine exposures. Indeed, we found only low frequencies of convergent B cell clones, and only expressed as unmutated IgM or IgD, in the cord blood repertoires (Fig. 1A, age zero years old). In contrast, convergent clones for Hib, NM, PP, TT, RSV, and Flu in children (one to three years of age) and adults (17-87 years of age) usually expressed mutated IgM or IgD or class-switched isotypes (Fig. 1A and fig. S1). There was a higher frequency of Hib, PP, TT and RSV convergent clones in the blood from children compared to adult blood (Fig. 1B). The childhood vaccination schedule (table S3) ensured immunization of the children to Hib, PP, and TT with immunizations at approximately 2, 4, 6 months, and again between 12-15 months of life; NM is not given as an early childhood vaccine, and showed low convergent clone frequencies (Fig. 1A). The timing of vaccination events compared to the detection of convergent IGH did not show significant correlation (fig. S2-S4), suggesting that children have persistently elevated frequencies of peripheral blood B cells expressing convergent clones for these pathogens.

**Fig. 1.**
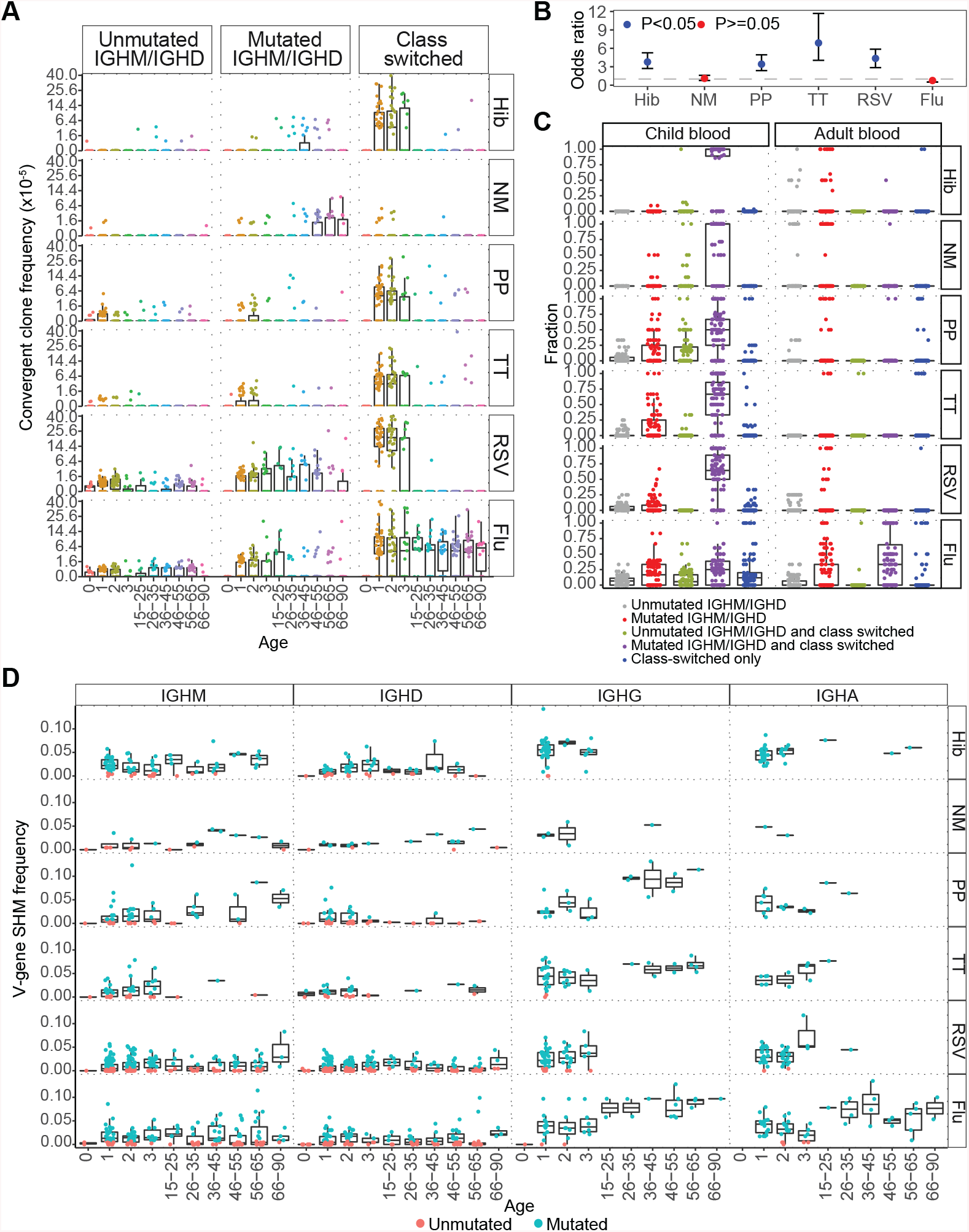
Frequency, class-switching and SHM state of pathogen-specific convergent clones in individuals of different ages. (A) The frequency of IGH sequences of convergent clones for different pathogens in blood samples from individuals of different ages. Each point represents the sum of clone frequencies for that pathogen in the indicated antibody type for that sample, with frequencies plotted on a square root scale. Convergent clones were split into unmutated IgM/IgD clones (**≤** 1% SHM), mutated IgM/IgD+ clones (>1% SHM) and class-switched clones. Ages are in years, with age 0 indicating cord blood. (B) Odds ratios (ORs) for detection of convergent clones in children compared to adults were calculated based on the fraction of convergent clones for each pathogen among the total clones in children and in adults. The points were colored by p-values calculated by Fisher’s exact test. (C) The fraction of convergent clones expressing unmutated IgM/IgD, mutated IgM/IgD, class-switched, or combinations of these three groups. Each point represents the fraction of convergent clones in each category to the total convergent clones for that pathogen in an individual. Cord blood samples are not shown, as almost all sequences in these specimens are unmutated IgM and IgD. To compare the frequency of class-switched convergent clones with mutated IgM/IgD clone members (colored in purple) between children and adults, the p-values were calculated by Wilcoxon–Mann-Whitney test, p-value = 5.08e-32, 6.66e-29, 8.25e-29, 9.93e-35, 1.71e-41 for Hib, NM, PP, TT, and RSV, respectively. The frequency of those convergent clones for Flu are not significantly different between children and adults. (D) The IGHV-gene SHM frequencies of each convergent clone in participants of different ages indicated in years. Each point represents the median IGHV-gene SHM frequency of each convergent cluster expressing each isotype. Clones with median SHM frequency less than or equal to 1% are colored in pink, and those with median SHM frequency more than 1% are in blue. The SHM frequencies of convergent clones expressing IgG or IgA were lower in children than in adults, p-value = 6.50e-13 and 1.96e-8, respectively, by Wilcoxon–Mann-Whitney test.

In analyzing antibody isotype expression by convergent clones, we classified clonal lineages into three groups: 1) those containing only unmutated IgM or IgD (unmutM/D), most likely derived from naïve B cells that have not undergone antigen-driven proliferation and diversification; 2) those containing IgM or IgD with median SHM frequencies over 1%, but no class-switched members (mutM/D); and 3) those containing class-switched members (CS) with or without IgM or IgD clone members. In adult blood, frequencies of mutM/D convergent clones specific for Hib, NM, and RSV were higher than unmutM/D convergent clones to these pathogens (fig. S5A). In contrast, the frequency of CS but not mutM/D convergent clones specific for PP, TT, and Flu in adults was higher than unmutM/D clones specific for these antigens (fig. S5A). These data suggest that non-switched memory B cell populations in adults are the predominant antigen-experienced B cells for some vaccines with prominent polysaccharide antigens (Hib, NM), but also for the glycoproteins of RSV. Surprisingly, higher frequencies of CS convergent clones were detected in child blood compared to adult blood for Hib, NM, PP, TT, and RSV (fig. S5B) even for pathogens with prominent polysaccharide antigens. A high fraction of the children’s CS convergent clones to Hib, NM, PP, TT, and RSV also contained mutated IgM expressing (IgM+) and IgD expressing (IgD+) clone members (Fig. 1C). Both adults and children had a high fraction of CS convergent clones for influenza (Fig. 1A).

Median SHM frequencies for IgM+ or IgD+ convergent clones for each pathogen were similar between adults and children (from 2% to 5%) whereas SHM frequencies of IgG+ or IgA+ convergent clones were significantly lower in children (Fig. 1D), in agreement with our prior observations of total IGH repertoires for these isotypes (*34*). We observed an increase of SHM up to age 87 years in IgG+ convergent clones for Flu, likely due to the recurrent exposures to influenza antigens by infection, or annual influenza vaccination, in contrast to the other pathogens.

Potential explanations for the lower frequencies of CS B cell convergent clones in adult compared to pediatric blood could be that the CS clone members gradually die off after childhood exposures, or that they are localized elsewhere in the body in adults, or that the clones specific for individual vaccines or pathogens are diluted by the larger overall pools of memory clones in adults and therefore at too low a frequency to be detected. To test the hypothesis that these cells were in other tissues in adults, we analyzed IGH repertoires in the spleen, mediastinal lymph node (MDLN), and mesenteric lymph node (MSLN) in eight adult deceased organ donors (table S1).

In each deceased organ donor, we compared overall clone distributions within and between secondary lymphoid tissues and to those found in blood. As expected, blood B cells had higher frequencies (mean proportion: 43.8%) of naïve-like clones expressing unmutated IgM or IgD, compared to lymphoid tissues (mean proportion: 20% for MDLN and MSLN, 18.7% for spleen) (fig. S6A). The blood showed limited sharing of clones with lymphoid tissues, while higher frequencies of shared clones were found among the lymph node and spleen, suggest larger clone sizes in the lymphoid tissues, and limited recirculation in the blood (fig. S6A). Although many clones were found in at least two tissues, the larger clones amongst those detected were not evenly distributed: each tissue was dominated by different clones (fig. S6B). Clonal lineages with members shared between different tissues fell into two main categories: those that express IgM and IgA, or those expressing IgG subclasses (fig. S7A, B). SHM frequencies positively correlated with the number of distinct tissue sites in which a clone was found (fig. S7C), suggesting that clones that have been stimulated the most by prior antigen exposures become the most widely distributed in the body.

The frequency of B cell convergent clones for Hib, NM, PP, TT, RSV, and Flu in lymph nodes and spleen of deceased organ donors was significantly higher than in blood (Fig. 2A). Prior clinical reports also suggest that different lymphoid tissues may have specialized functions in protective immunity, such as enrichment of B cells specific for bacterial capsular polysaccharides in splenic tissue, leading to vulnerability to these pathogens in splenectomized patients (*28–33*). We tested whether each tissue site showed tissue-specific convergent IGH to particular pathogens, for example, whether spleen had more clonal lineages specific for polysaccharide antigens. Surprisingly, the frequencies of convergent clones for Hib, NM and PP in the two lymph node sites were similar or even higher than in spleen, indicating that the spleen may be a major reservoir of such clones, but is not the only source of them. Convergent IGH for polysaccharide antigens were usually expressed as IgM or IgD, and less often IgG (Fig. 2B) in lymphoid tissues. Convergent clones for PP were found expressing IgG in MSLN and spleen, and IgA in MDLN, however, indicating that the convergent clones for polysaccharide antigens can also be class-switched. One of the deceased organ donors (Donor 5) had a cause of death reported as anoxia related to influenza H1N1 infection with the 2009 pandemic strain. We noted that that this deceased donor had very few convergent clones for influenza in any of their lymphoid tissues or blood (fig. S8). Although a single example does not permit generalization, we had anticipated that the infection would have stimulated expansion of convergent clones for influenza; it seems possible that the lack of such clones was associated with an ineffective immune response to the infection.

**Fig. 2.**
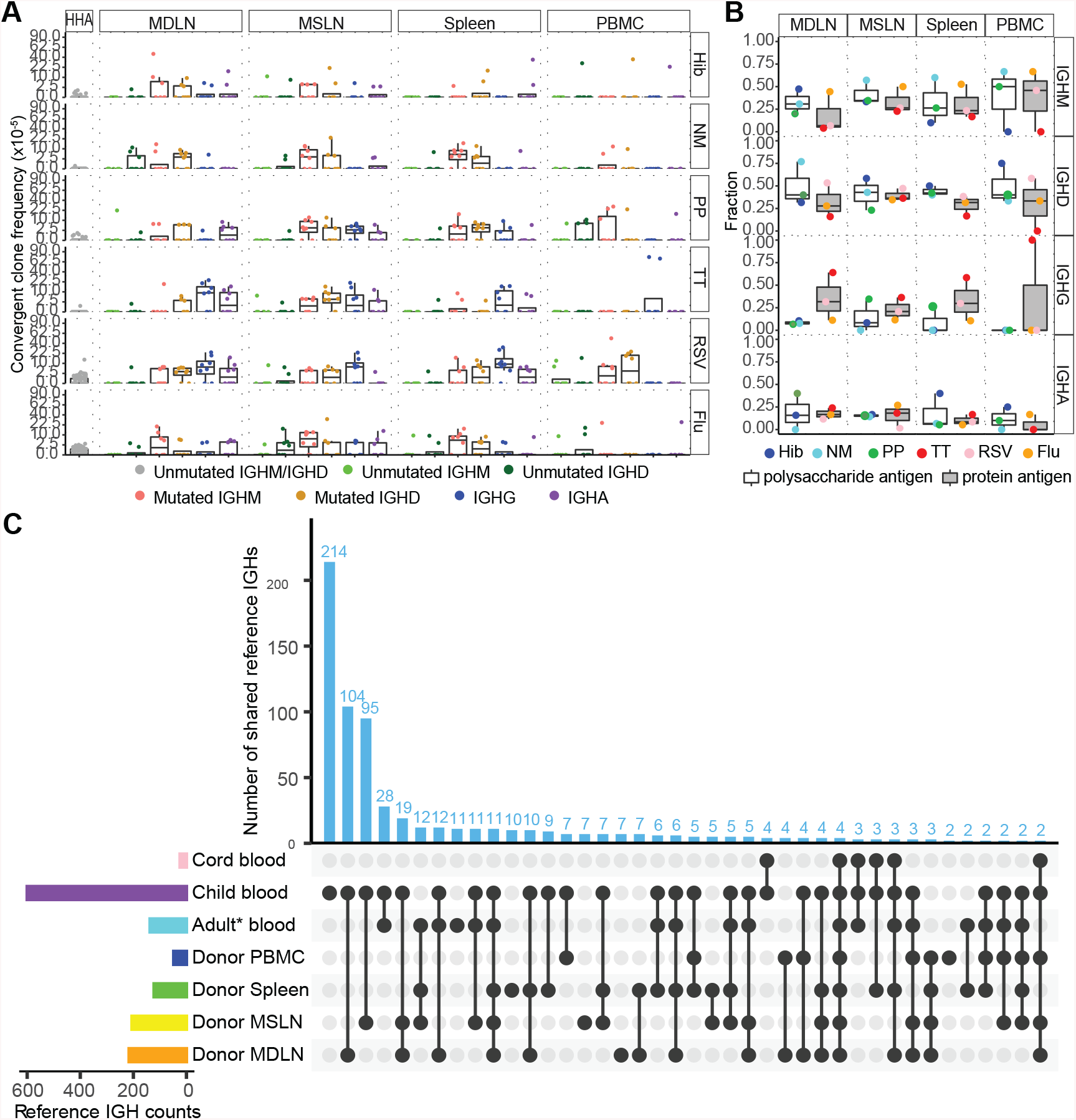
Convergent B cell clone distribution in tissues. (A) Frequency of unmutated IgM or IgD convergent B cell clones for each pathogen in 114 healthy adult (HHA) blood samples (left) and frequency of convergent B cell clones for each pathogen in the blood (PBMC), mediastinal lymph node (MDLN), mesenteric lymph node (MSLN) and spleen of eight deceased organ donors. Points are colored by isotype and plotted on a square root scale. The frequencies of convergent clones in lymphoid tissues were higher than in the blood, p-value = 0.00049, 0.0037, 0.016, 6.71e-7, 0.012, 0.00017 for Hib, NM, PP, TT, RSV, and Flu respectively, by Wilcoxon– Mann-Whitney test. (B) The fraction of convergent clones expressing each isotype in each tissue of the deceased organ donors. The points are colored by antigen and divided into polysaccharide (white box) and protein antigens (grey box). Convergent clones for polysaccharide antigens were more likely to express IgM and IgD (p-value = 0.035) and less likely to express IgG (p-value = 0.0058), p-values were calculated by Wilcoxon–Mann-Whitney test. (C) The distribution of shared known antigen-specific IGH sequences among cord blood, child blood, adult blood, and lymphoid tissues of the donors (spleen, MSLN, MDLN). The adult blood samples from the eight deceased organ donors were labeled as Donor PBMC, blood samples from other healthy adults are labeled as Adult* blood. The number of reference antigen-specific IGH sequences shared by each combination of specimen types is indicated by the vertical bars, with the dots and connecting lines in the lower part of the plot indicating the combination of specimen types. The total number of convergent IGHs found in a particular specimen type is indicated in the bars to the left of the plot. Child blood and the two lymph nodes of deceased adult organ donors had the highest numbers of shared known pathogen-specific IGH sequences, p-value = 0.0001181, by Fisher’s exact test.

We hypothesized that some differences in humoral immune responses in children compared to adults could be related to changes in antigen-specific B cell repertoires with development and aging. To test this, we evaluated whether distinct subsets of the reference IGH sequences specific for each pathogen or vaccine were enriched in particular age groups. In fact, IGH sequences from the two lymph nodes of the deceased organ donors, and the blood from the children were more likely to match the same known pathogen-specific IGH sequences, compared to IGH sequences from adult blood (including both the eight donor blood and the other 114 adult blood samples), indicating that localization of these clones differs between children and adults (Fig. 2C and fig. S9). Mean CDR-H3 lengths (fig. S10A) and IGHV gene usage (fig. S10, B and C) of convergent clones were not significantly different between cord blood, pediatric or adult blood, or adult lymphoid tissues.

Prompted by a recent serological report of cross-reactive SARS-CoV-2 binding antibodies in the blood of children prior to the COVID-19 pandemic (*24*), we next tested if we could detect convergent B cell clones for pathogens that had not been encountered by the study subjects. It has been reported that 40%-60% of healthy people with no history of SARS-CoV-2 infection mount CD4+ T-cell responses to peptides from this virus (*36*), but published evidence for SARS-CoV-2-binding B cells in unexposed people has been limited to relatively rare individuals who had prior infection with SARS-CoV (*37*), or examples likely derived from naive B cells expressing stereotypic IGH with IGHV3-53/IGHV3-66 and IGHJ6 gene segments (*38*). The infants, children and adults in this study had not been exposed to SARS-CoV-2 or Ebola virus (EBOV). We detected rare examples of convergent clones for EBOV in the unmutM/D compartment in both adult and pediatric blood, and in adult lymphoid tissues (Fig. 3, A and B). In stark contrast, convergent clones with IGH similar to SARS-CoV-2 binding and neutralizing antibodies (Table 1, table S4) were common in the blood of children one to three years old, and in 37 of 51 children showed SHM with or without class-switching, providing evidence of prior antigen exposure (Fig. 3A, C). Cord blood lacked these mutated and class-switched sequences, further supporting a role for antigen stimulation in their development (Fig. 3A). Adults had lower frequencies of SARS-CoV-2 convergent clones in blood compared to the children, with some IgM or IgD clones having SHM, but almost no class-switched examples (Fig. 3A). We tested whether convergent SARS-CoV-2 B cell clones in adults might be localized to lymphoid tissues rather than circulating in the blood, but found similarly low clone frequencies, with comparable SHM frequencies to those in the blood, without class switching, in lymph nodes and spleen (Fig. 3B, C, D). Notably, about 7% of the SARS-CoV-2 convergent IGHs in children were similar to known antibodies that cross-react with other human coronaviruses (HCoVs), such as HKU1, NL63, and 229E, which are continuously circulating in the population (*37*) (Fig. 3E). None of the convergent clones for SARS-CoV-2 in adults were similar to IGH previously reported to bind other HCoVs, with the exception of SARS-CoV. These results suggest that children have higher frequencies than adults of cross-reactive spike-binding memory B cells stimulated by other HCoV exposures, leading to the pre-pandemic cross-reactive SARS-CoV-2 binding and neutralizing antibody titers recently described in children(*24*). We note that both adults and children had relatively high frequencies of convergent SARS-CoV-2 antibody clones detected in naive B cell-derived unmutated IgM and IgD, indicating that these clonotypes have high generation frequencies in the primary B cell repertoires (Fig. 3A). The cross-reactive B cell memory and serum antibodies could contribute to the milder illnesses caused by SARS-CoV-2 in children, in combination with other factors such as decreased expression of the angiotensin-converting enzyme 2 (ACE2) viral receptor protein in the airway (*39*). Relatively rapid decreases in antibody titers specific for various coronaviruses in adults have been previously reported (*40, 41*), suggesting a lack of long-lived plasma cells producing secreted antibodies. Given that the adults in our study were once children, and are likely to have been exposed periodically to community coronaviruses through their lifetimes, another implication of our data is that coronavirus infections may not stimulate long-lasting memory B cell responses, particularly of class-switched B cells.

**Fig. 3.**
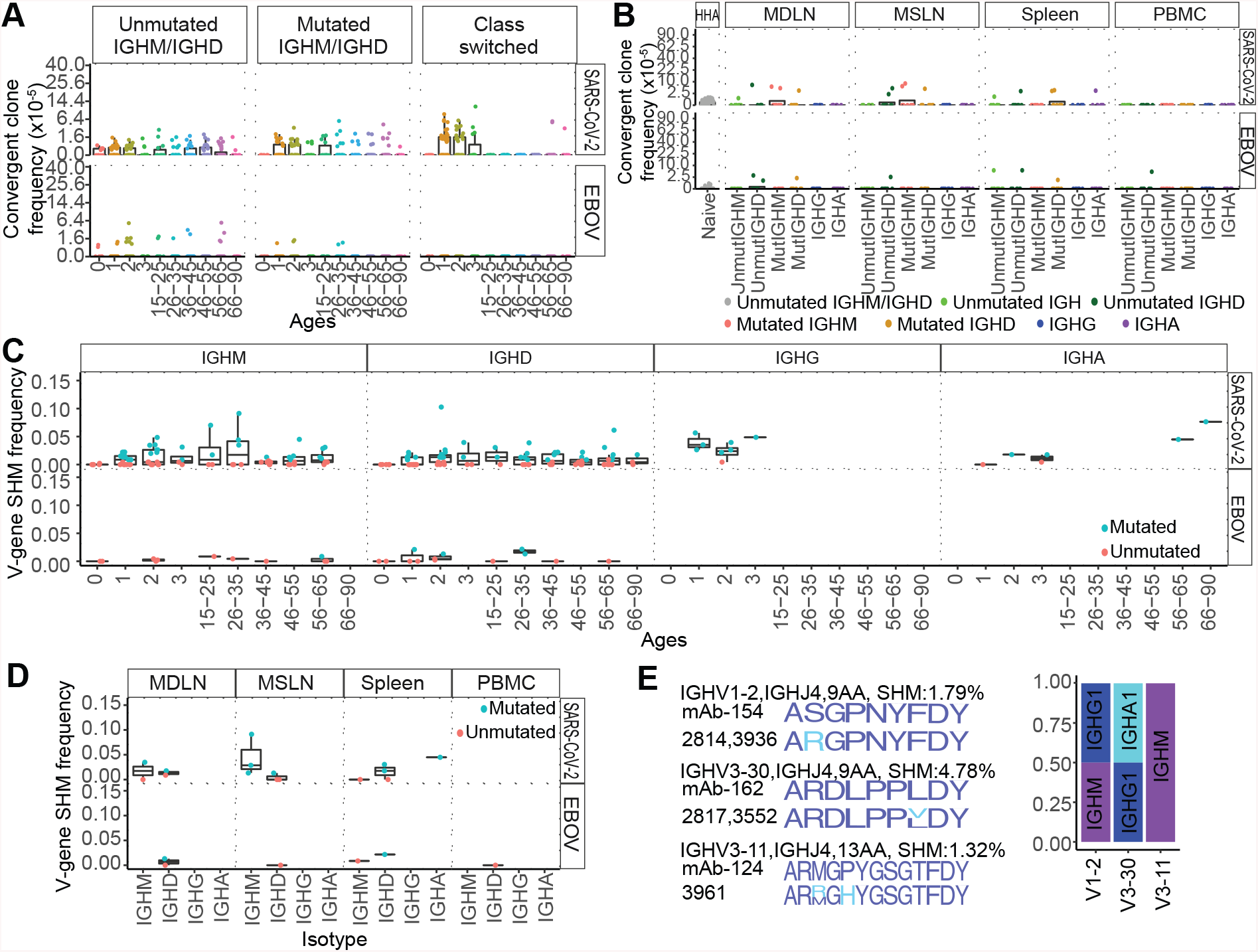
Convergent clones for SARS-CoV-2 and EBOV. (A) The frequency of convergent clones for SARS-CoV-2 and EBOV in individuals of different ages. Convergent clones were split into unmutated IgM/IgD clones (SHM **≤** 1%), mutated IgM/IgD+ clones (SHM >1%) and isotype switched clones, with clone frequencies plotted on a square root scale. For SARS-CoV-2, children had significantly higher frequency of mutated convergent clones with or without class-switched isotypes compared to unmutated IgM/IgD clones (p-value = 0.00012, by Wilcoxon– Mann-Whitney test). The frequency of class-switched convergent clones and the frequency of mutated IgM/IgD convergent clones for SARS-CoV-2 was higher in children than in adults (p-values = 1.22e-13 and 0.0089, respectively, by Wilcoxon–Mann-Whitney test). (B) The frequency of convergent clones for SARS-CoV-2 and EBOV in the blood and lymphoid tissues of deceased organ donors. Points are colored by isotype and SHM level. (C) The SHM frequencies of convergent clones expressing a given isotype in participants of different ages (x-axis). Clones with median SHM frequency less than or equal to 1% are colored in pink, while those with median SHM frequency greater than 1% are in blue. (D) The SHM frequencies of convergent clone members of each isotype in lymphoid tissue or blood of deceased organ donors. Clones with median SHM frequency less than or equal to 1% are colored in pink, while those with median SHM frequency greater than 1% are in blue. (E) Sequence logos of CDR-H3 amino acid residues of convergent IGH cross-reactive to SARS-CoV-2 and other human coronaviruses. For each of the three sets of convergent IGH, the sequence logo for the reported antigen-specific CDR-H3 was shown in the top row; sequence logos for clones from children were aligned below (colored blue where they match the reported CDR-H3, colored cyan if different). To the right the isotype proportion of convergent clones similar to each SARS-CoV-2-specific IGH is shown. The isotype proportion is calculated based on the number of reads expressing each isotype divided by the total number of reads in each convergent clone. The median SHM frequency for each convergent clone is shown after the convergent clone label.

**Table 1.**
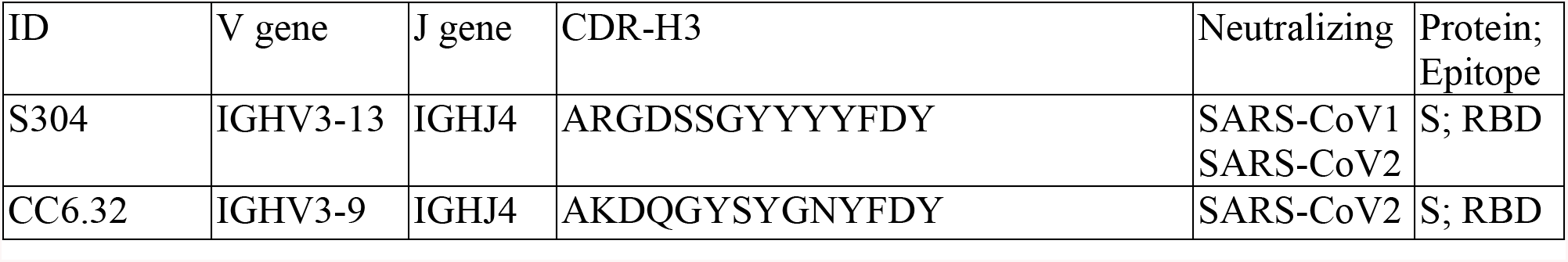
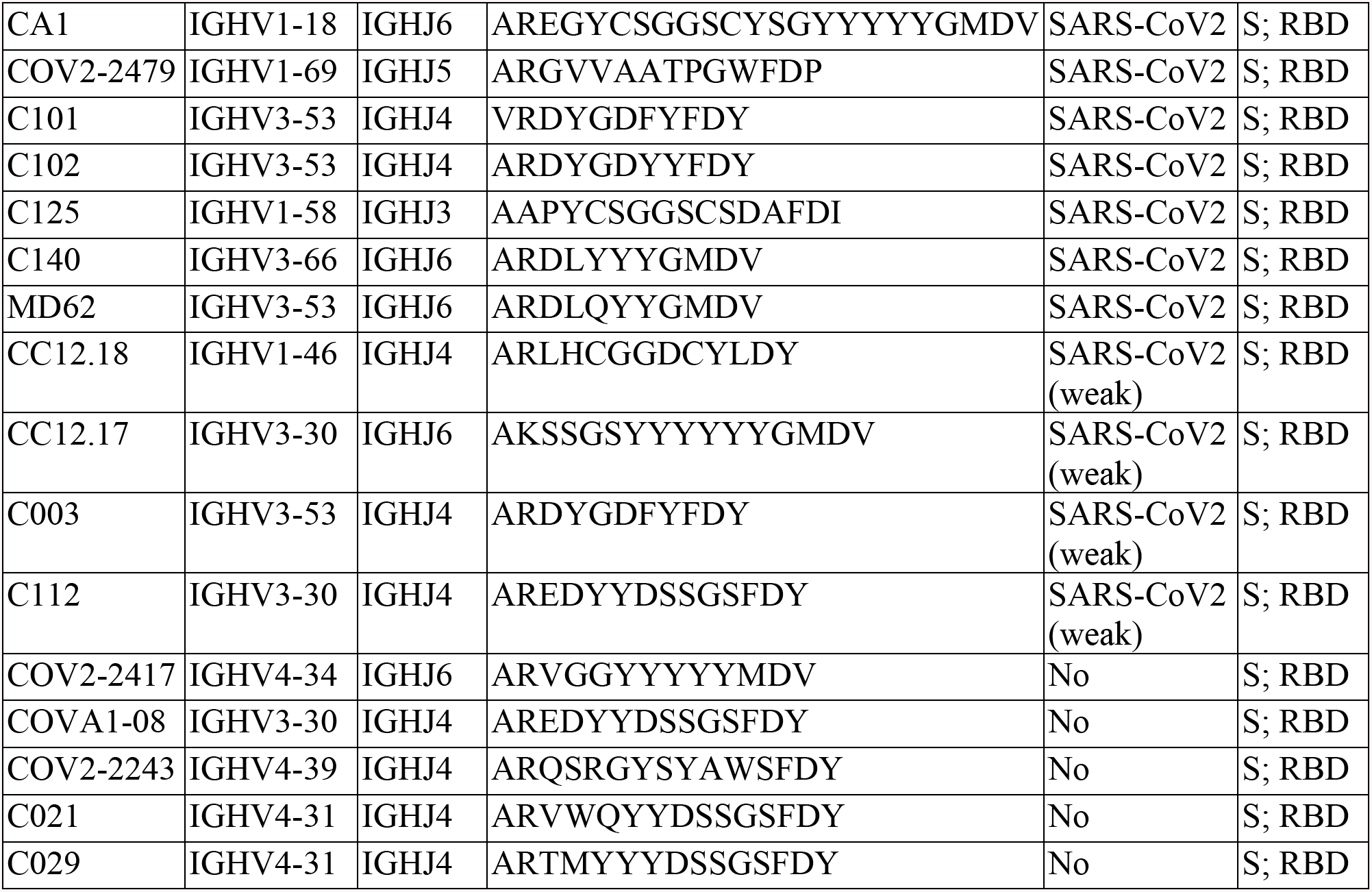
List of reported SARS-CoV-2 antibodies that can be found in child blood samples and which are reported as neutralizing antibodies or can bind to the RBD of SARS-CoV-2.

Our findings clarify several features of human B cell selection in people of different ages, and in different lymphoid tissues in adults. The immune responses of early childhood are particularly important in an individual’s life, as they form the original memory B cell pools and shape future responses (*42*). Our data show that individuals with diverse underlying genotypes can respond to common pathogens and vaccine antigens in a conserved manner (producing some highly similar convergent clones in response to the same antigen), and those convergent clones are found at higher frequency and more often as class-switched isotypes in the blood of children compared to adults. Among adults, these convergent clones are instead enriched in lymph nodes and spleen. B cell memory to antigens that are frequently encountered by infection or vaccination, such as those of influenza viruses, show similar clone frequencies and isotype switching between children and adults, but demonstrate progressive increases in SHM frequencies over the human lifespan. In contrast to ideas of tissue-enriched B cell populations for particular antigens, such as bacterial capsular polysaccharide-specific B cells in the spleen, the B cell clones likely to be specific for these antigens are found at similar frequencies in the spleen and lymph nodes.

Our data also highlight important differences in potential cross-reactive B cell responses to novel pathogens in children and adults. Young children and adults both showed very little evidence for IGH specific for EBOV, consistent with their lack of exposure to Ebola or related viruses. Notably, young children have class-switched and mutated clones with similarity to known SARS-CoV-2 specific IGH, particularly for clones known to be cross-reactive for binding the spike antigens of SARS-CoV-2 and other coronaviruses, raising the possibility that previous HCoV exposures provide preexisting immunity to SARS-CoV-2 in children. The lack of such class-switched cross-reactive memory in adults, in both blood and lymphoid tissues, adds further support for the notion that serum antibody responses as well as B cell memory to coronaviruses may be short-lived. More detailed knowledge of the role of pre-existing cross-reactive memory B cell populations in new immune responses, and the determinants of long-lived B cell memory and plasma cell formation will be important for the development of safe, effective vaccines to SARS-CoV-2 as well as other pathogens.

## Acknowledgments

We thank all staff members of the California Transplant Donor Network (now Donor Network West). Special thanks to Sharon Swain, RN, MSN. We thank Dr. Andrew Z. Fire, Dr. Elyse A. Hope and Dr. Jason D. Merker for helpful discussions and contributions to the research.

## Author contributions

F.Y. and S.D.B. conceived of the project; F.Y., S.C.A.N., K.J.L.J., and K.M.R performed data analyses; F.Y., S.C.A.N., and S.D.B. verified the analyses; R.A.H., R.S.O., E.L.M., J-Y.L., and T.D.P. contributed to sample preparation and carried out the experiments; C.U.N., and S.D.B. supervised and supported the project; F.Y., S.C.A.N., and S.D.B. wrote the initial manuscript; and all authors provided critical feedback and contributed to the final manuscript;

## Data and materials availability

All data is available in the main text or the supplementary materials. Previously generated IGH repertoire data are available with BioProject number PRJNA503602 (child data set) and PRJNA491287 (114 healthy human adult data set reported by Nielsen et al. (*34*)). The IGH repertoire data for the deceased organ donors and the cord blood infant samples is in BioProject: PRJNA674610.

## References and Notes

1. M. E. Buckingham, S. M. Meilhac, Tracing cells for tracking cell lineage and clonal behavior. Dev Cell 21, 394–409 (2011).

2. T. W. LeBien, T. F. Tedder, B lymphocytes: how they develop and function. Blood 112, 1570–1580 (2008).

3. E. V. Rothenberg, Immune Cell Identity: Perspective from a Palimpsest. Perspect Biol Med 58, 205–228 (2015).

4. D. Jung, C. Giallourakis, R. Mostoslavsky, F. W. Alt, Mechanism and control of V(D)J recombination at the immunoglobulin heavy chain locus. Annu Rev Immunol 24, 541–570 (2006).

5. D. D. Dudley, J. Chaudhuri, C. H. Bassing, F. W. Alt, Mechanism and control of V(D)J recombination versus class switch recombination: similarities and differences. Adv Immunol 86, 43–112 (2005).

6. C. W. Davis et al., Longitudinal Analysis of the Human B Cell Response to Ebola Virus Infection. Cell 177, 1566–1582 e1517 (2019).

7. K. R. McCarthy, D. D. Raymond, K. T. Do, A. G. Schmidt, S. C. Harrison, Affinity maturation in a human humoral response to influenza hemagglutinin. Proc Natl Acad Sci U S A, (2019).

8. I. Setliff et al., Multi-Donor Longitudinal Antibody Repertoire Sequencing Reveals the Existence of Public Antibody Clonotypes in HIV-1 Infection. Cell Host Microbe 23, 845–854 e846 (2018).

9. A. Watanabe et al., Antibodies to a Conserved Influenza Head Interface Epitope Protect by an IgG Subtype-Dependent Mechanism. Cell 177, 1124–1135 e1116 (2019).

10. S. F. Andrews et al., Preferential induction of cross-group influenza A hemagglutinin stem-specific memory B cells after H7N9 immunization in humans. Sci Immunol 2, (2017).

11. K. J. Jackson et al., Human responses to influenza vaccination show seroconversion signatures and convergent antibody rearrangements. Cell Host Microbe 16, 105–114 (2014).

12. P. Parameswaran et al., Convergent antibody signatures in human dengue. Cell Host Microbe 13, 691–700 (2013).

13. C. J. Henry Dunand, P. C. Wilson, Restricted, canonical, stereotyped and convergent immunoglobulin responses. Philos Trans R Soc Lond B Biol Sci 370, (2015).

14. J. Truck et al., Identification of antigen-specific B cell receptor sequences using public repertoire analysis. J Immunol 194, 252–261 (2015).

15. A. K. Simon, G. A. Hollander, A. McMichael, Evolution of the immune system in humans from infancy to old age. Proc Biol Sci 282, 20143085 (2015).

16. L. Goo, V. Chohan, R. Nduati, J. Overbaugh, Early development of broadly neutralizing antibodies in HIV-1-infected infants. Nat Med 20, 655–658 (2014).

17. F. Gotzinger et al., COVID-19 in children and adolescents in Europe: a multinational, multicentre cohort study. Lancet Child Adolesc Health 4, 653–661 (2020).

18. X. Lu et al., SARS-CoV-2 Infection in Children. N Engl J Med 382, 1663–1665 (2020).

19. F. Zheng et al., Clinical Characteristics of Children with Coronavirus Disease 2019 in Hubei, China. Curr Med Sci 40, 275–280 (2020).

20. J. Cai et al., A Case Series of children with 2019 novel coronavirus infection: clinical and epidemiological features. Clin Infect Dis, (2020).

21. N. Parri, M. Lenge, D. Buonsenso, G. Coronavirus Infection in Pediatric Emergency Departments Research, Children with Covid-19 in Pediatric Emergency Departments in Italy. N Engl J Med 383, 187–190 (2020).

22. I. Liguoro et al., SARS-COV-2 infection in children and newborns: a systematic review. Eur J Pediatr 179, 1029–1046 (2020).

23. R. Carsetti et al., The immune system of children: the key to understanding SARS-CoV-2 susceptibility? Lancet Child Adolesc Health 4, 414–416 (2020).

24. K. W. Ng et al., Preexisting and de novo humoral immunity to SARS-CoV-2 in humans. Science, (2020).

25. S. P. Weisberg et al., Distinct antibody responses to SARS-CoV-2 in children and adults across the COVID-19 clinical spectrum. Nature Immunology, (2020).

26. C. Berek, C. Milstein, Mutation drift and repertoire shift in the maturation of the immune response. Immunol Rev 96, 23–41 (1987).

27. J. Jacob, G. Kelsoe, K. Rajewsky, U. Weiss, Intraclonal generation of antibody mutants in germinal centres. Nature 354, 389–392 (1991).

28. C. G. Vinuesa, C. de Lucas, M. C. Cook, Clinical implications of the specialised B cell response to polysaccharide encapsulated pathogens. Postgrad Med J 77, 562–569 (2001).

29. W. Timens, A. Boes, T. Rozeboom-Uiterwijk, S. Poppema, Immaturity of the human splenic marginal zone in infancy. Possible contribution to the deficient infant immune response. J Immunol 143, 3200–3206 (1989).

30. A. Cerutti, M. Cols, I. Puga, Marginal zone B cells: virtues of innate-like antibody-producing lymphocytes. Nat Rev Immunol 13, 118–132 (2013).

31. I. C. MacLennan, Y. J. Liu, Marginal zone B cells respond both to polysaccharide antigens and protein antigens. Res Immunol 142, 346–351 (1991).

32. A. J. van den Eertwegh, J. D. Laman, M. M. Schellekens, W. J. Boersma, E. Claassen, Complement-mediated follicular localization of T-independent type-2 antigens: the role of marginal zone macrophages revisited. Eur J Immunol 22, 719–726 (1992).

33. G. Leone, E. Pizzigallo, Bacterial Infections Following Splenectomy for Malignant and Nonmalignant Hematologic Diseases. Mediterr J Hematol Infect Dis 7, e2015057 (2015).

34. S. C. A. Nielsen et al., Shaping of infant B cell receptor repertoires by environmental factors and infectious disease. Sci Transl Med 11, (2019).

35. M. Bracht, D. Basevitz, M. Cranis, R. Paulley, Impact of respiratory syncytial virus: the nurse’s perspective. Drugs R D 11, 215–226 (2011).

36. A. Grifoni et al., Targets of T Cell Responses to SARS-CoV-2 Coronavirus in Humans with COVID-19 Disease and Unexposed Individuals. Cell 181, 1489–1501 e1415 (2020).

37. A. Z. Wec et al., Broad neutralization of SARS-related viruses by human monoclonal antibodies. Science 369, 731–736 (2020).

38. S. I. Kim et al., Stereotypic Neutralizing V<sub>H</sub> Clonotypes Against SARS-CoV-2 RBD in COVID-19 Patients and the Healthy Population. bioRxiv, 2020.2006.2026.174557 (2020).

39. S. Bunyavanich, A. Do, A. Vicencio, Nasal Gene Expression of Angiotensin-Converting Enzyme 2 in Children and Adults. JAMA 323, 2427–2429 (2020).

40. K. A. Callow, H. F. Parry, M. Sergeant, D. A. Tyrrell, The time course of the immune response to experimental coronavirus infection of man. Epidemiol Infect 105, 435–446 (1990).

41. A. W. D. Edridge et al., Seasonal coronavirus protective immunity is short-lasting. Nat Med, (2020).

42. A. Olin et al., Stereotypic Immune System Development in Newborn Children. Cell 174, 1277–1292 e1214 (2018).

