## Supplementary material for "Shared B cell memory to coronaviruses and other pathogens varies in human age groups and tissues": Supplemenntal File

### Materials and Methods

Tissue collection protocol

Approval from the Committee on Human Research at the University of California San Francisco for collecting deceased (brain dead) organ donor specimens was not required because deceased donors are not considered human subjects under Federal law. As governed by the Uniform Anatomical Gift Act (UAGA), all deceased organ donors had documentation of separate authorization for donation and research, respectively. Authorization was provided either as first-person authorization (for example, registration with the Department of Motor Vehicles, DMV) or legal next-of-kin. As such, The Research Committee of the California Transplant Donor Network (now Donor Network West) approved the collection of specimens from deceased organ donors who had research authorization. All deceased donors were clinically managed according to established protocols for organ donation, including administration of 5g methylprednisolone. Whole blood samples and three lymphoid tissue types: spleen, mediastinal lymph nodes, and mesenteric lymph nodes were collected during the organ recovery process. Lymphoid tissue samples were rapidly frozen on dry ice. Whole blood samples were collected in standard serum gold-top tubes and placed in an ice-filled biohazard bag. Lymphoid tissue specimens were frozen in liquid nitrogen following collection. Deceased organ donors received steroids for hemodynamic stabilization prior to the collection of organs for transplant; while we cannot exclude the possibility that this protocol could perturb the distribution of B cells in the tissues and potentially cause cell death, it is less likely that it would alter the IGHV gene usage, SHM level and isotype expression in individual B cells.

Cord blood sample collection

Cord blood samples were collected in potassium EDTA anticoagulated blood collection tubes and stored in 4°C, with volumes ranging from 2-5 ml. After 7 days of storage at 4°C, unopened cord blood specimens not used for further clinical diagnostics were retrieved (Stanford IRB protocol 3124). Samples were processed through density gradient separation using Histopaque-1077 (Sigma-Aldrich), and peripheral blood mononuclear cells (PBMCs) were isolated. PBMCs were aliquoted, flash-frozen, and stored in -80°C until needed for library preparation.

PCR amplification of IGH libraries for high-throughput DNA sequencing (HTS)

AllPrep column purification (Qiagen, Valencia, CA) was used to isolate RNA. Complementary DNA (cDNA) was generated from total RNA using SuperScript™ III (Invitrogen) and random hexamer priming (Promega Corporation, C1181). PCR amplification of IGH rearrangements from the cDNA template for HTS on the Illumina MiSeq instrument was carried out according to published protocols (*1*). Each template was amplified using multiplexed IGHV primers based on the BIOMED-2 primer set in the framework one region and one isotype-specific primer located in the first exon of the constant region (*1*). These first round PCR primers also encoded approximately half of the Illumina adapter sequences. The first round PCR used AmpliTaq Gold (Applied Biosystems) enzyme, with final primer concentrations of 3.3 μM, and the following program: 94°C for 7 min, 35 cycles of (94°C for 30 sec, 58°C for 45 sec, 72°C for 120 sec), and a final extension at 72°C for 10 min. Illumina adapters were completed by a second PCR carried out with the Qiagen Multiplex PCR kit (Qiagen), using 0.4 μL of the first PCR product as the template in a 30 μL reaction with the following program: 94°C for 15 min, 12 cycles of (94°C for 30 sec, 60°C for 45 sec, 72°C for 90 sec), and a final extension at 72°C for 10 min. Primer sequences for all libraries are in table S5. Each isotype was amplified separately to decrease chimeric product generation. PCR reactions for all samples were pooled and purified by agarose gel electrophoresis and gel extracted using the QIAquick kit (Qiagen). Libraries were sequenced as a PE300 run using a v3 600-cycle kit.

<u>IGH read processing and analysis</u>

IGH reads were merged and separated by barcode (exact match) followed by primer trimming. V, D, J gene segments, and junctional bases were assigned using a local installation of the IgBLAST program (*2*). Isotype and subclass were determined by aligning non-primer-encoded constant region sequences to IMGT database reference sequences requiring an exact match (*3*). Somatic mutations in the IGHV-region were identified by alignment with gapped IMGT germline reference IGHV sequences. To avoid attributing PCR or sequencing error sequence changes to SHM, we chose a conservative threshold of 1% difference from the germline IGHV sequence to call a sequence mutated (*1, 4*).

Annotation of antibody isotype and subtype of RNA-derived sequences was accomplished by looking for perfect matches between constant regions from the IMGT (*5*) database and sequence upstream of the constant region primer. CDR-H3 sequences were identified on the basis of the conserved cysteine-104 and the motif downstream of the conserved tryptophan-118 residue 37. Non-IGH artifactual sequences and those with poor IGHV matches (IGHV-segment match bit score less than 140) were removed from the data. The python scripts and R code used for secondary analyses of the data can be provided on request.

In total, 13,385,447 sequences from cord blood samples (mean: 1,115,453, median: 1,089,792 per individual); 68,874,117 sequences from child blood (mean: 1,250,472, median: 1,193,687 per individual; and mean:740,581, median: 761,661 per sample); 68,831,446 sequences from healthy controls (mean: 603,785, median: 637,269 per individual) were obtained and analyzed. For the deceased organ donor cohort, at total of 23,238,446 sequences from the blood or each lymphoid tissue site (mean: 580,961, median:558,572 per sample) were obtained and analyzed.

All data from this study are deposited in the SRA database, in BioProjects: PRJNA674610 (deceased organ donor and infant cord blood data), PRJNA503602 (child blood data set) and PRJNA491287 (114 healthy human adult data set reported by Nielsen et al. (*6*)).

IGH reference sequences

Antigen-specific IGH sequences specific for different antigens were collected from different publications (table S2), including polysaccharide vaccines targeting three bacterial pathogens, 1) *Haemophilus influenzae* type b (Hib) (*7-10*), 2) *Neisseria meningitidis* (NM) (*11-13*), and 3) *Pneumococcus pneumoniae* (PP) (*14-17*); IGH sequences specific for protein antigens 1) tetanus toxoid (TT) vaccine (*18-21*), 2) respiratory syncytial virus (RSV) fusion (F) protein (*22-26*), 3) influenza virus (*27-42*), 4) SARS-CoV-2 (*43-51*), and 5) Ebola virus (*52*). The polysaccharide vaccines targeting pneumococcus reported by Chen *et al.* (*14*) and Adler *et al.* (*17*) are conjugated to Diphtheria CRM197 protein.

Inference of clonal lineages for IGH sequences amplified from cDNA

To control for variability in amplification and abundance of RNA-derived sequences and to follow a clonal lineage across different isotype and mutation states, sequences were clustered into putative clonal lineages using single-linkage clustering (python package scipy.cluster.hierarchy (*53*)) as follows: reads from the same individual were clustered if they shared the same IGHV segment, IGHJ segment (not taking into account the allele calls), and CDR-H3 length, as well as a CDR-H3 nucleotide sequence identity of at least 90%. The identification of members of particular clones using the single-linkage clustering method can vary somewhat with sequencing depth in a dataset, but the conclusions obtained in this analysis were robust with regard to sequencing depth.

Calculation of convergent clones for each pathogen

To identify convergent rearranged IGH among different individuals for each pathogen, reference antigen-specific IGH sequences and IGH sequences from individuals in our study annotated with the same IGHV and IGHJ segment (not considering alleles) and the same CDR-H3 length were clustered based on 85% CDR-H3 amino acid sequence similarity. Partitioning of sequences by V, J and CDR-H3 length was performed using custom software, while the CDR-H3 amino acid sequence clustering was performed using CD-HIT with options -c 0.85 -l 4 -S 0 -g 1 -b 1 (*54*). Clusters were selected as informative if they contained at least two IGH sequences from each individual and matched to at least one reference antigen-specific IGH sequence.

Statistical analysis

Analysis of IGH sequence data was performed with custom python code using pandas and NumPy (*55*). Statistical tests were performed in R (*56*) using base packages for statistical analysis and the ggplot2 package for graphics. Box-whisker plots show median (horizontal line), interquartile range (box), and 1.5 times the interquartile range (whiskers). P-values were calculated by using Wilcoxon-Mann-Whitney test, Fisher's exact test or one-way ANOVA with Tukey's HSD test. The statistical methods and p-values for each analysis are provided in the results or figure legends.

### Supplementary Figures

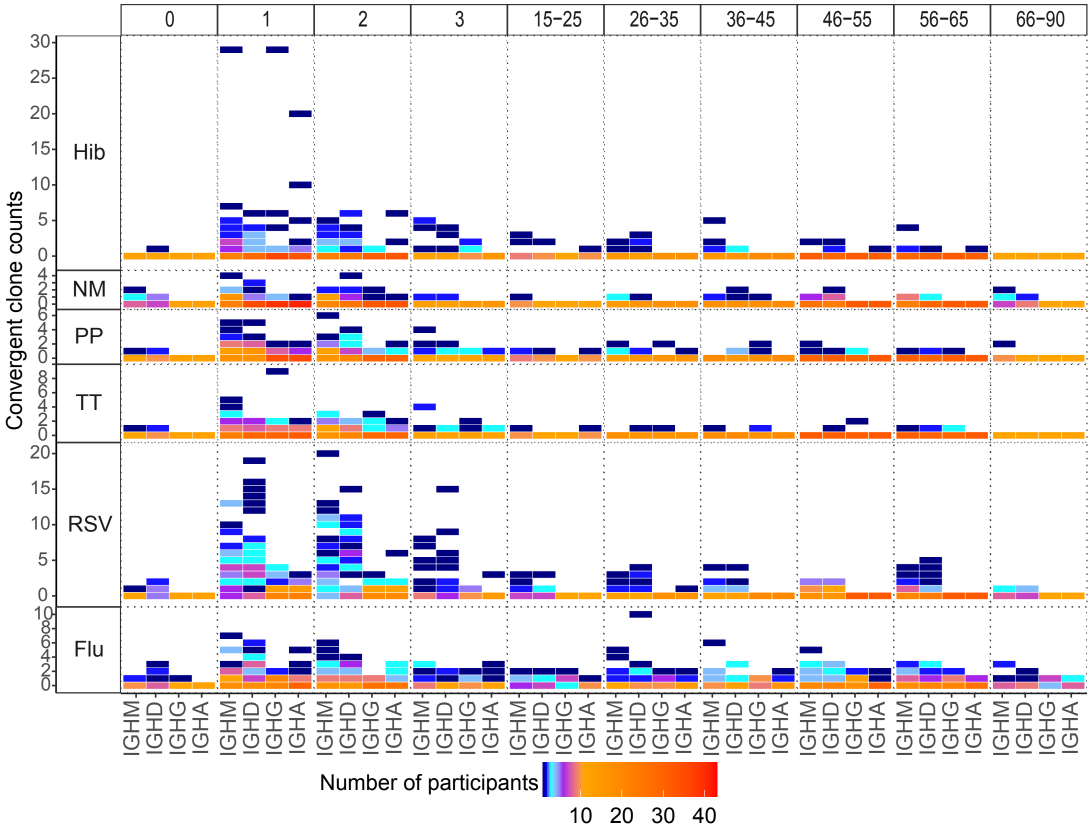


**Fig. S1. Number of participants with given number of convergent clones for each pathogen in blood at different ages.** Heat map showing the number of samples with a given number of convergent clones for each pathogen in blood of individuals with different ages. The x-axis is the isotype, and the y-axis is the number of convergent clones, and the color represents the number of participants. Panel columns indicate age and panel rows show pathogen.

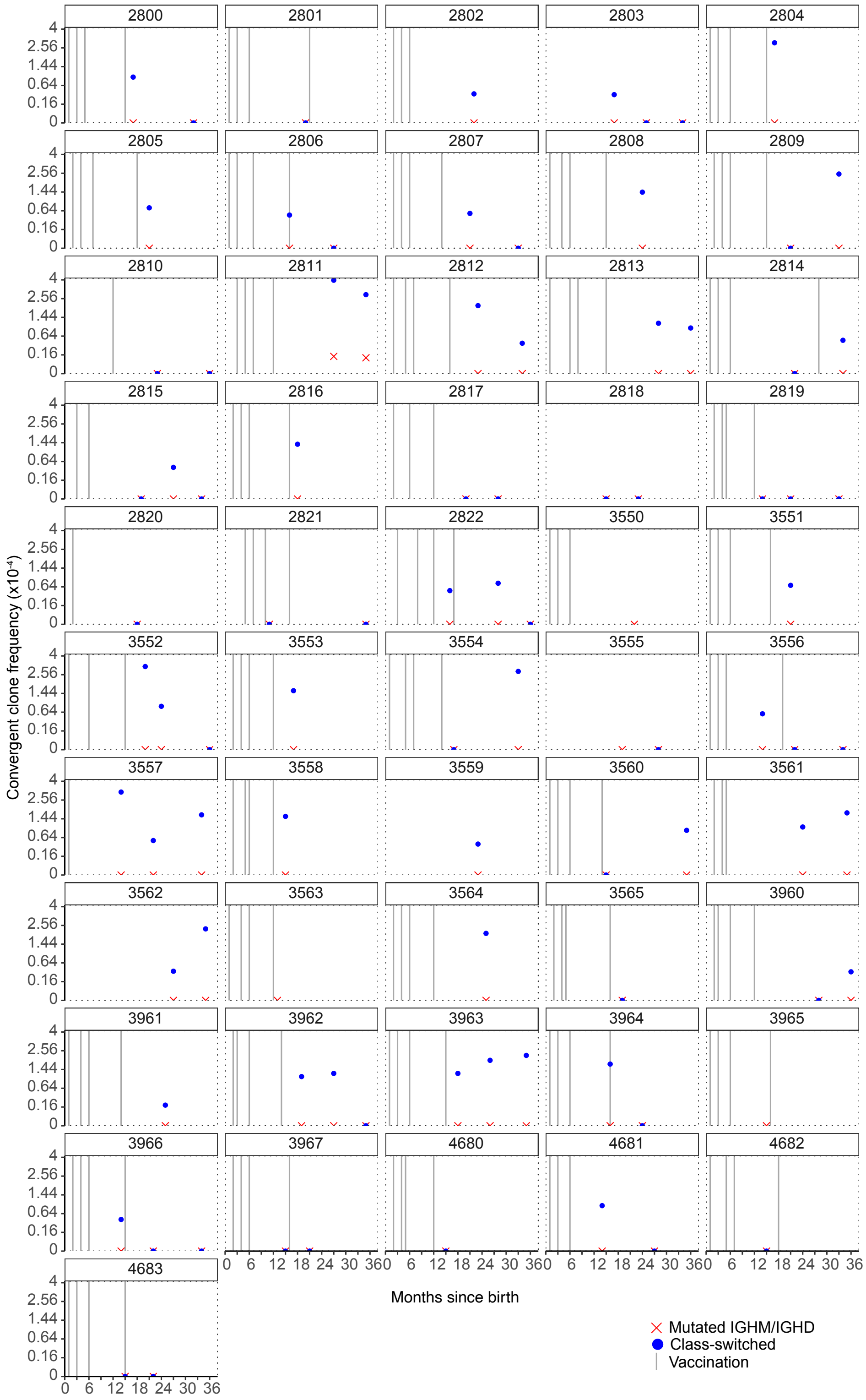


**Fig. S2. Frequency of convergent clone for Hib and vaccination histories of children.** The frequency of mutM/D convergent clones was shown as red crosses, the frequency of CS convergent clones is shown as blue dots, and the time of vaccinations is indicated by the grey vertical lines. For some children, the dates of vaccination were unknown. In each child, the frequency of convergent clones for each pathogen in the mutM/D or CS sequence pools was calculated by dividing the total number of convergent clones for that pathogen in each sample by the total number of clones of that group (mutM/D or CS) in the sample.

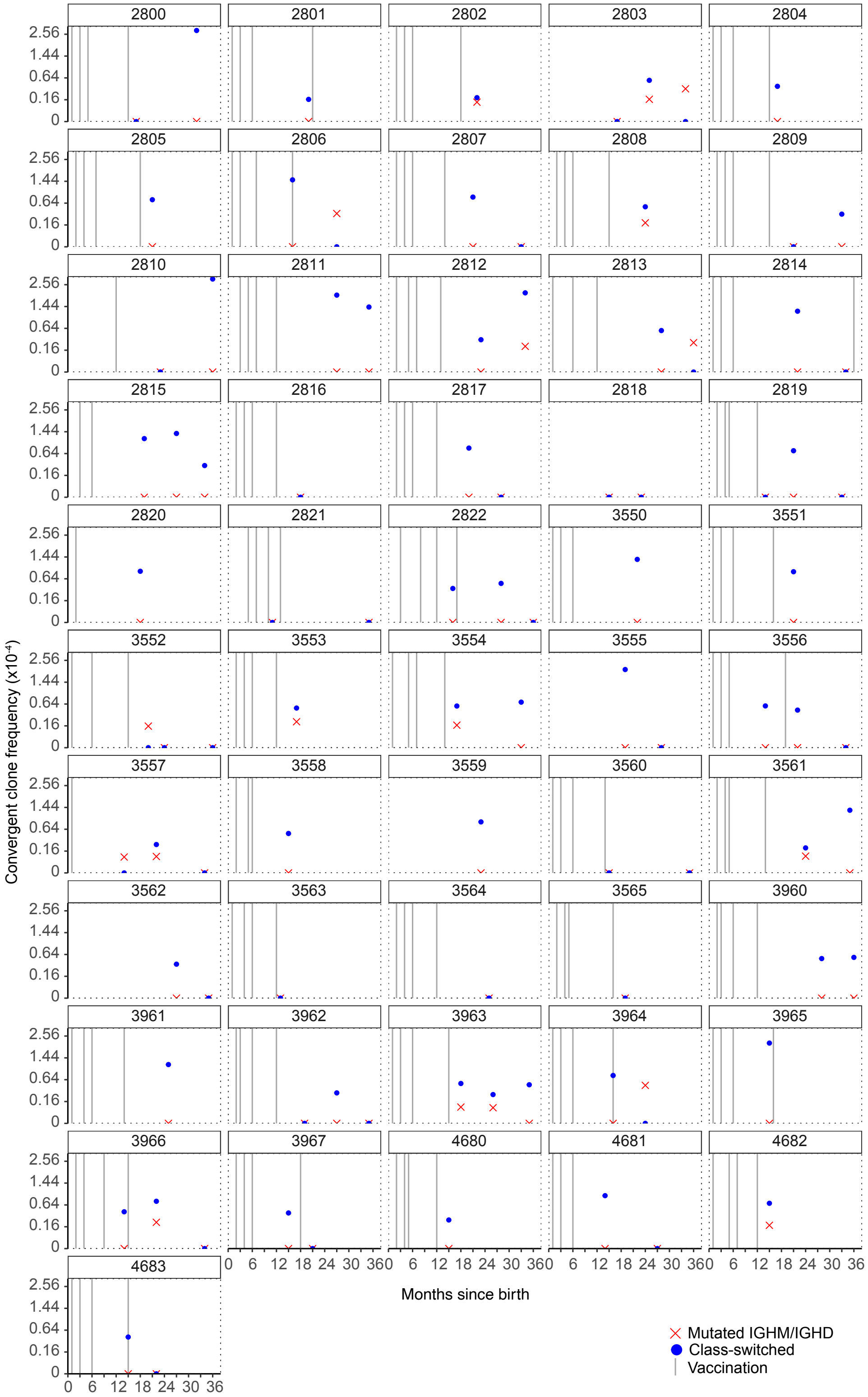


**Fig. S3. Frequency of convergent clones for PP and vaccination histories of children.** The frequency of mutM/D convergent clones was shown as red crosses, the frequency of CS convergent clones is shown as blue dots and the time of vaccinations was shown as grey vertical lines. For some children, the dates of vaccination were unknown. In each child, the frequency of convergent clones for each pathogen in the mutM/D or CS sequence pools was calculated by dividing the total number of convergent clones for that pathogen in each sample by the total number of clones of that group (mutM/D or CS) in the sample.

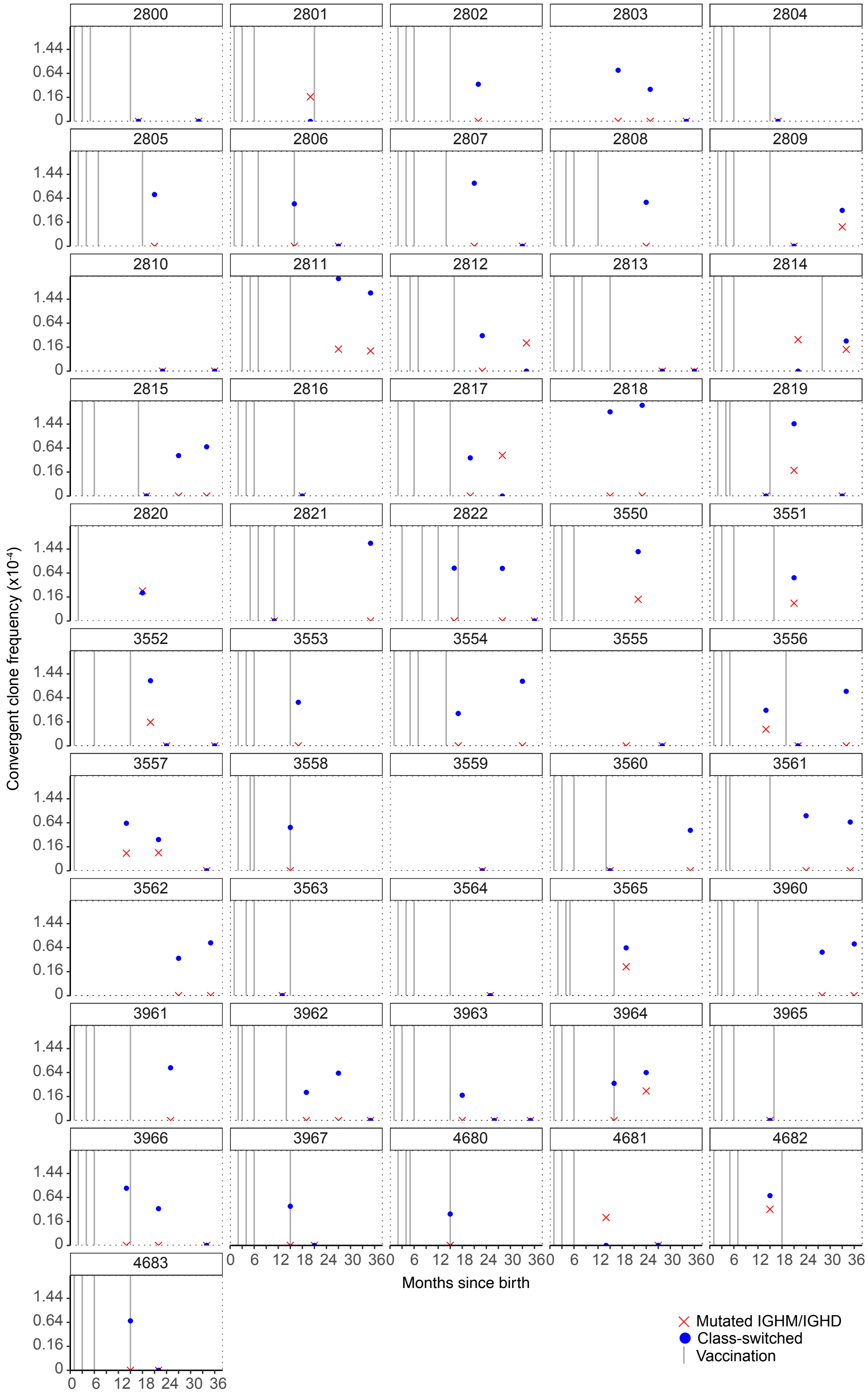


**Fig. S4. Frequency of convergent clones for TT and vaccination histories of children.** The frequency of mutM/D convergent clones was shown as red crosses, the frequency of CS convergent clones was shown as blue dots and the time of vaccinations was shown as grey vertical lines. For some children, the dates of vaccination were unknown. In each child, the frequency of convergent clones for each pathogen in the mutM/D or CS sequence pools was calculated by dividing the total number of convergent clones for that pathogen in each sample by the total number of clones of that group (mutM/D or CS) in the sample.

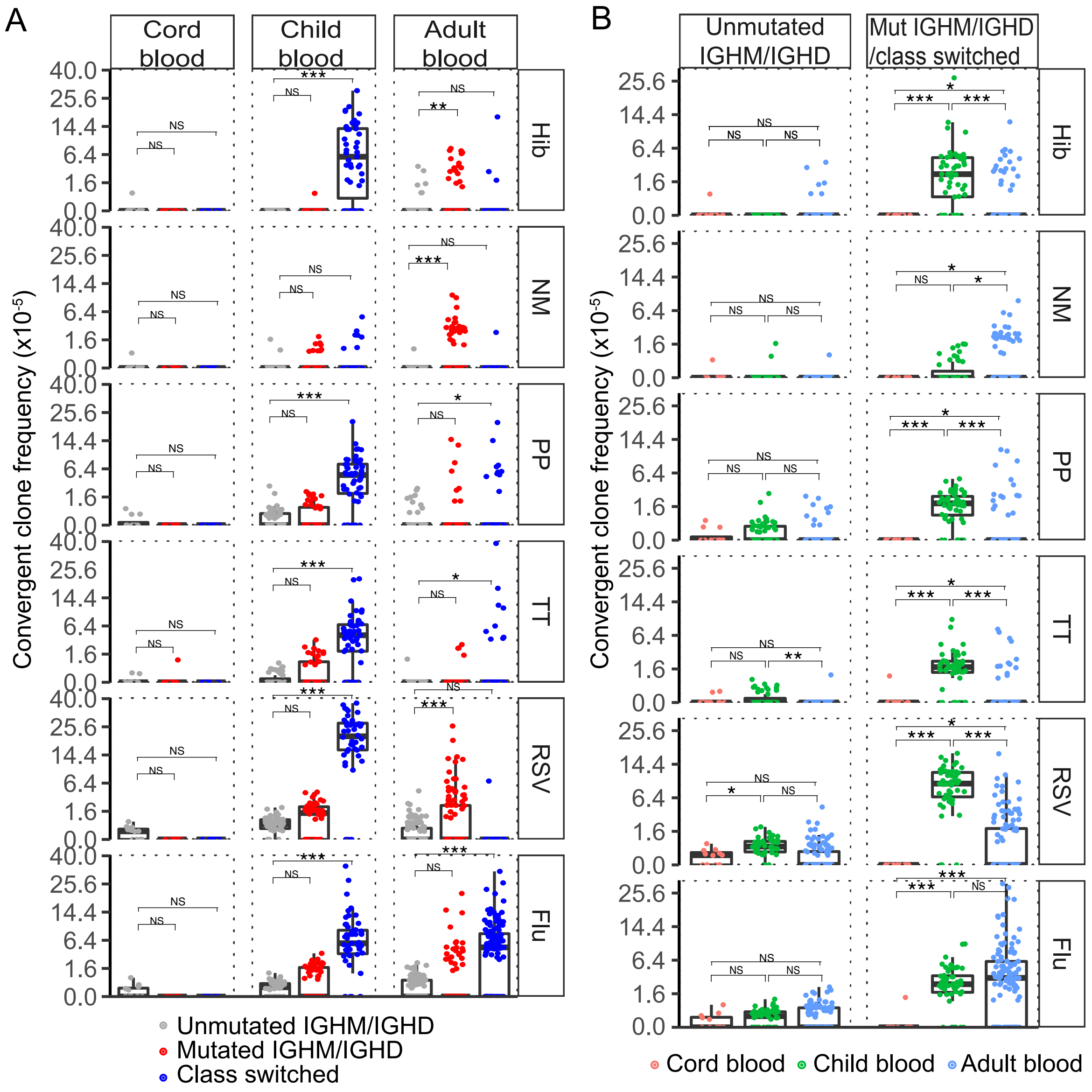


**Fig. S5. Frequencies of different convergent clone types for each pathogen in individuals of different ages.** (A) The frequency of the three convergent clone types for different pathogens in individuals of different ages. Each dot represents the clone frequency within an individual, plotted on a squared root scale. (B) The frequency of convergent clones in participants of different age groups. Convergent clones were split into unmutated IgM/IgD convergent clones (≤1% V gene SHM) or convergent clones with either mutated IgM/IgD or class-switched members. Each dot represents the frequency within an individual and plotted at the squared root scale. For (A) and (B), the convergent clone frequency is calculated by the total number of convergent clones of each group for each pathogen divided by the total number of clones of the corresponding group in each individual. The differences between groups were tested using one-way ANOVA with Tukey's HSD test. ***p-value < 0.001; **p-value < 0.01; *p-value ≤ 0.05; NS: p-value > 0.05. (A) and (B) are plotted using the same samples but are plotted in different ways to highlight different features of the responses.

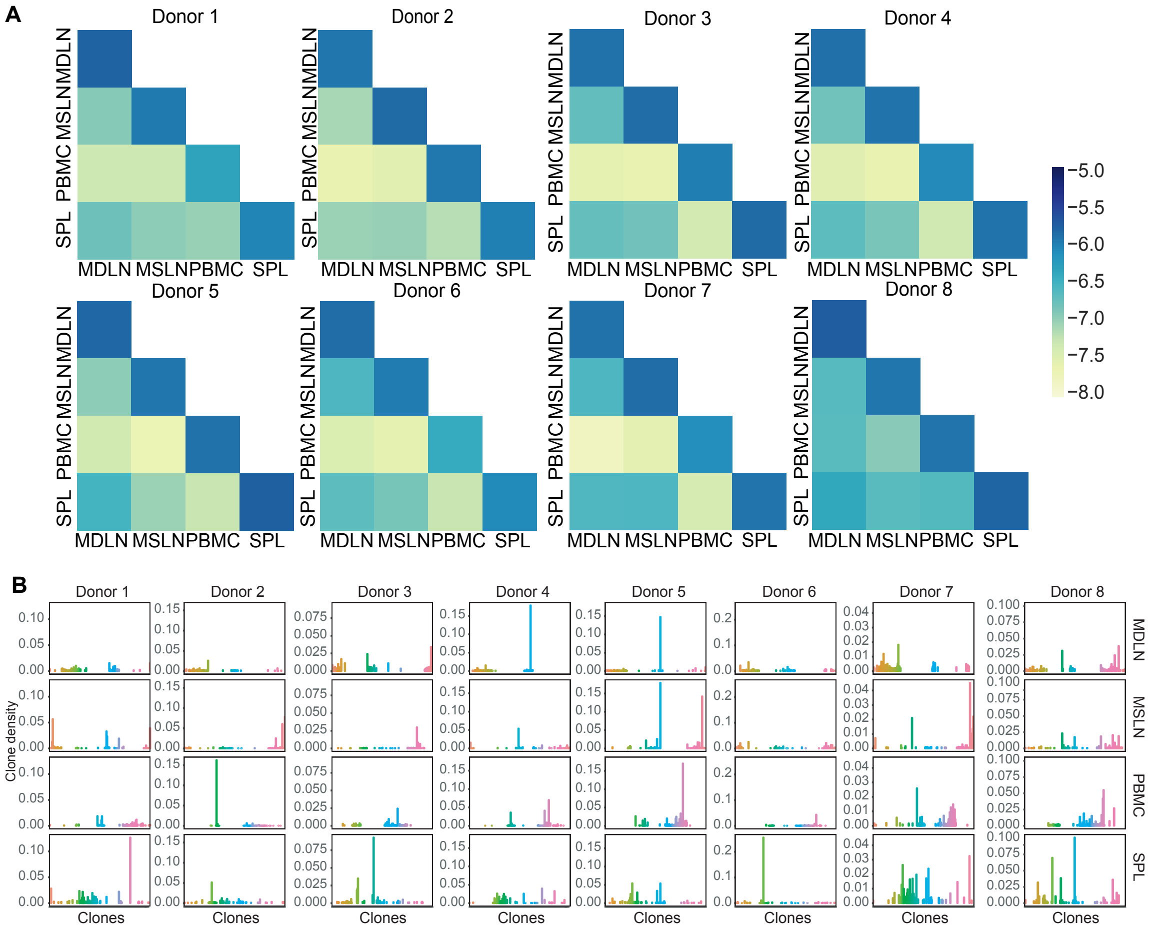


**Fig. S6. Distribution of B cell clones in different tissues of deceased organ donors.** (A) Heat map of B cell clonal sharing between each specific lymphoid tissue type of each deceased organ donor. The diagonals for each tissue represent the probability that two randomly selected reads belong to the same clone. The off diagonals represent the probability that a randomly selected read from one tissue and a randomly selected read from the second tissue are members of the same clone. The filling color scale is the negative log scale of the clone counts controlled by the product of reads in the corresponding two samples. Compared to the clone sharing between lymphoid tissues and blood, higher frequencies of shared clones were found among the lymph node and spleen, p-value = 3.285e-12 by Wilcoxon–Mann-Whitney test. (B) The density of the top 50 largest clones in each tissue of each deceased organ donor. Bar plot showing the frequency of reads from each B cell clone in the same order for each tissue type. An arbitrary color was assigned to each clone. Only the 50 highest frequency clones are displayed. SPL: Spleen

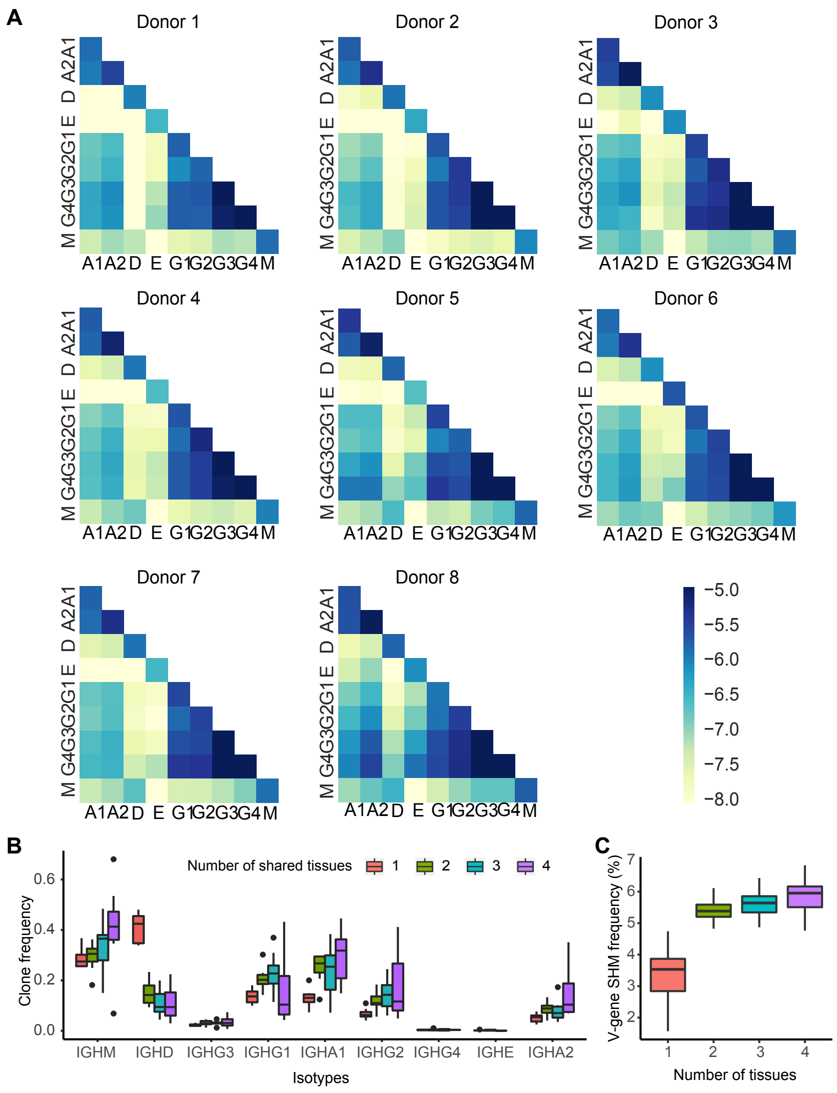


**Fig. S7. Distribution and SHM frequency of B cell clones expressing different isotypes in deceased organ donors.** (A) Heat map of B cell clonal sharing between B cells expressing different isotypes/sub-isotypes of each donor. The diagonals represent the probability that two randomly selected reads of the same isotype/sub-isotype are members of the same clone. The off diagonals represent the probability that a randomly selected read from one isotype/sub-isotype and a randomly selected read from the second isotype/sub-isotype are members of the same clone. The color scale is the negative log scale of the clone counts controlled by the product of reads in the corresponding two samples. M:IGHM, D:IGHD, G1:IGHG1, G2:IGHG2,

G3:IGHG3, G4:IGHG4, A1:IGHA1, A2:IGHA2, E:IGHE (B) The frequency of B cell clones expressing different isotypes occurring in one, two, three, or four studied tissues. The boxes in the box-whisker plots were colored by the number of shared tissues. (C) SHM frequency of B cell clones detected in one, two, three, or four tissues in each donor. The y-axis is the mean SHM frequency. Box plot color encodes the number of tissues in which the B cell clone was detected.

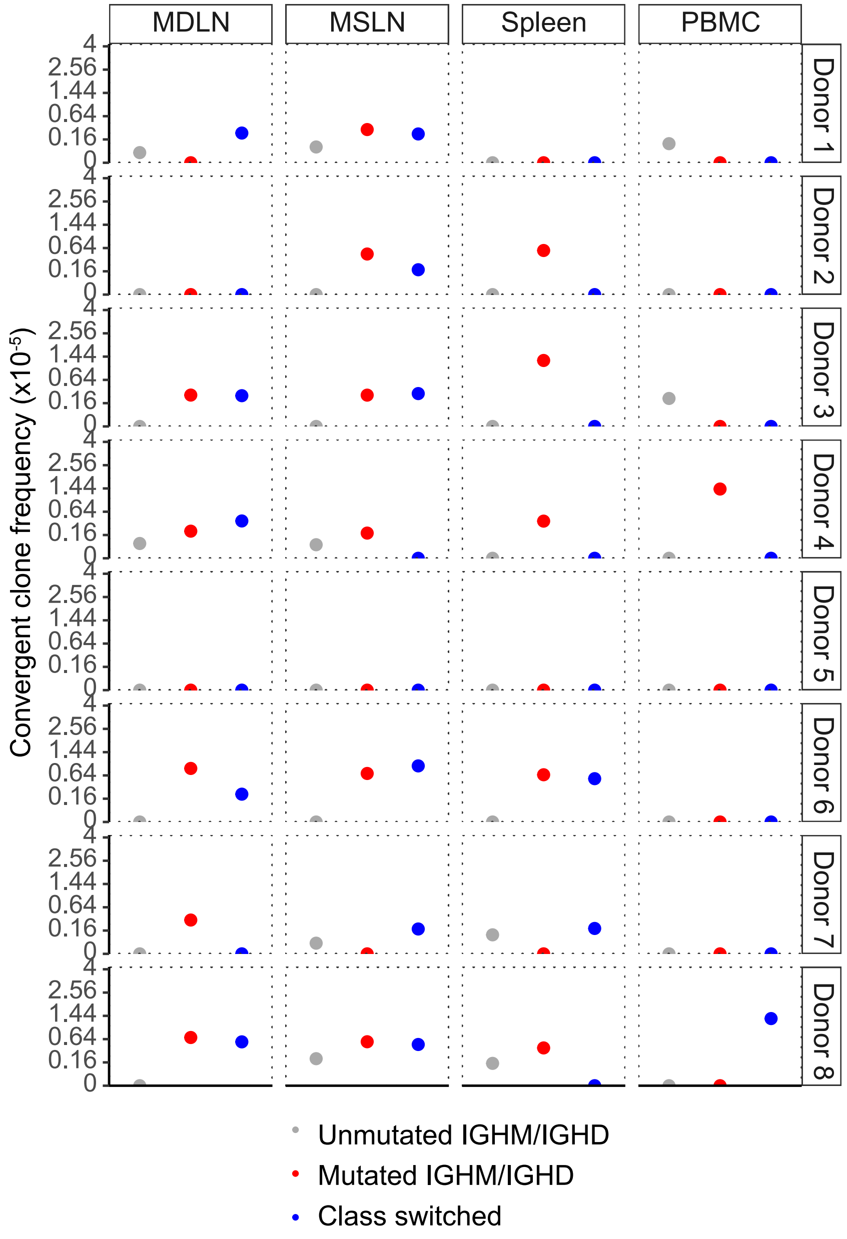


**Fig. S8. Frequency of convergent clones for influenza in lymphoid tissues and blood of deceased organ donors.** The frequency of convergent clones for flu in deceased organ donors are plotted on a squared root scale. UnmutM/D convergent clones are shown as dark grey dots, mutM/D convergent clones are shown as red dots, and CS convergent clones are show as blue dots. In each sample, the frequency of convergent clones for each pathogen for mutM/D or CS was calculated by dividing the total number of convergent clones of a given group by the total number of clones of the corresponding group.

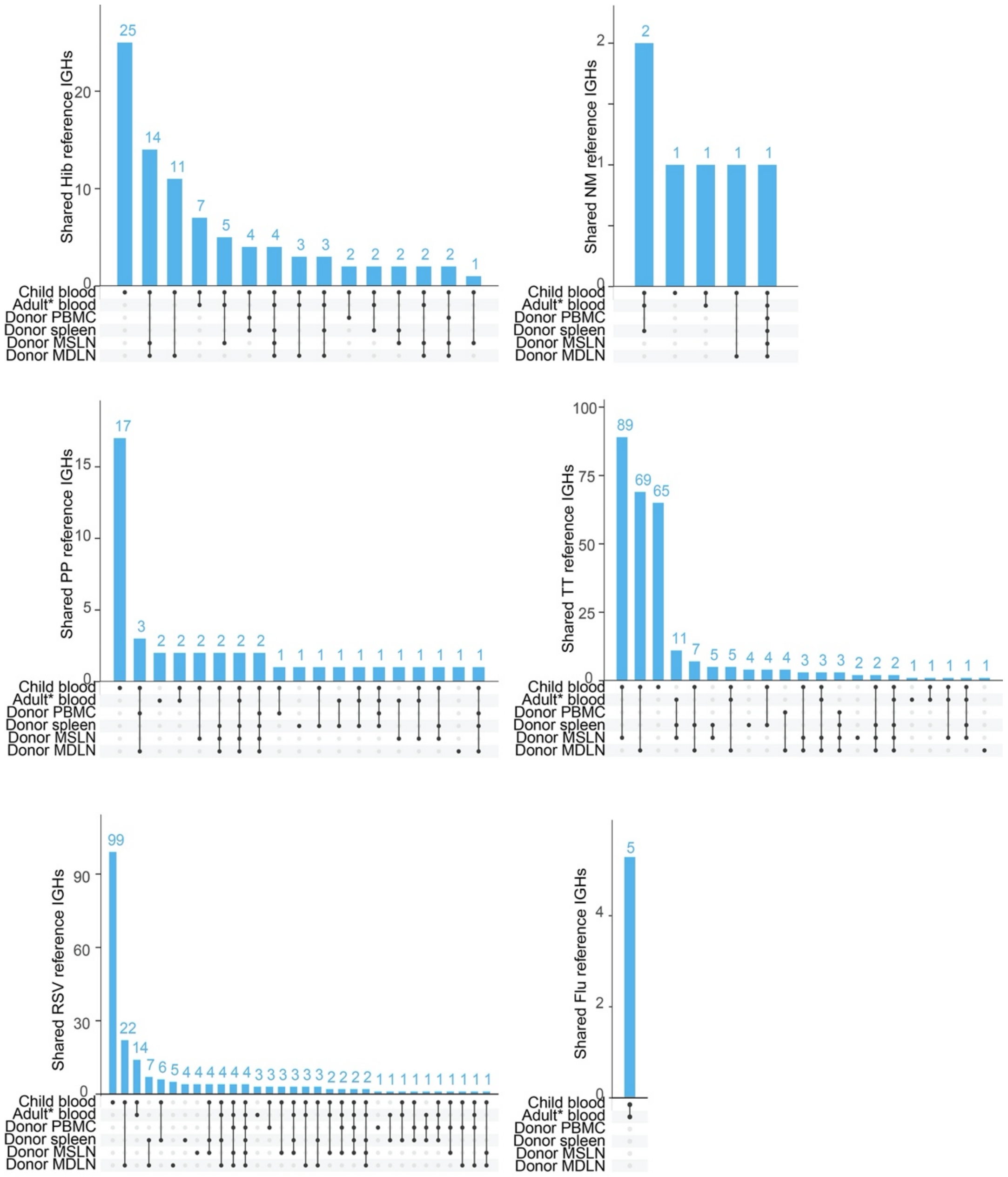


**Fig. S9. Convergent IGH sequences shared by children's blood, adult blood, and adult tissues.** The number of known antigen-specific IGH shared by children, adult blood and different lymphoid tissues of adults. The adult blood samples from the eight deceased organ donors were labeled as Donor PBMC, blood samples from other healthy adults are labeled as Adult* blood. The number of reference antigen-specific IGH sequences shared by each combination of specimen types is indicated by the vertical bars, with the dots and connecting lines in the lower part of the plot indicating the combination of specimen types. The total number of convergent IGHs for each pathogen found in a particular specimen type is indicated in the bars to the left of the plot.

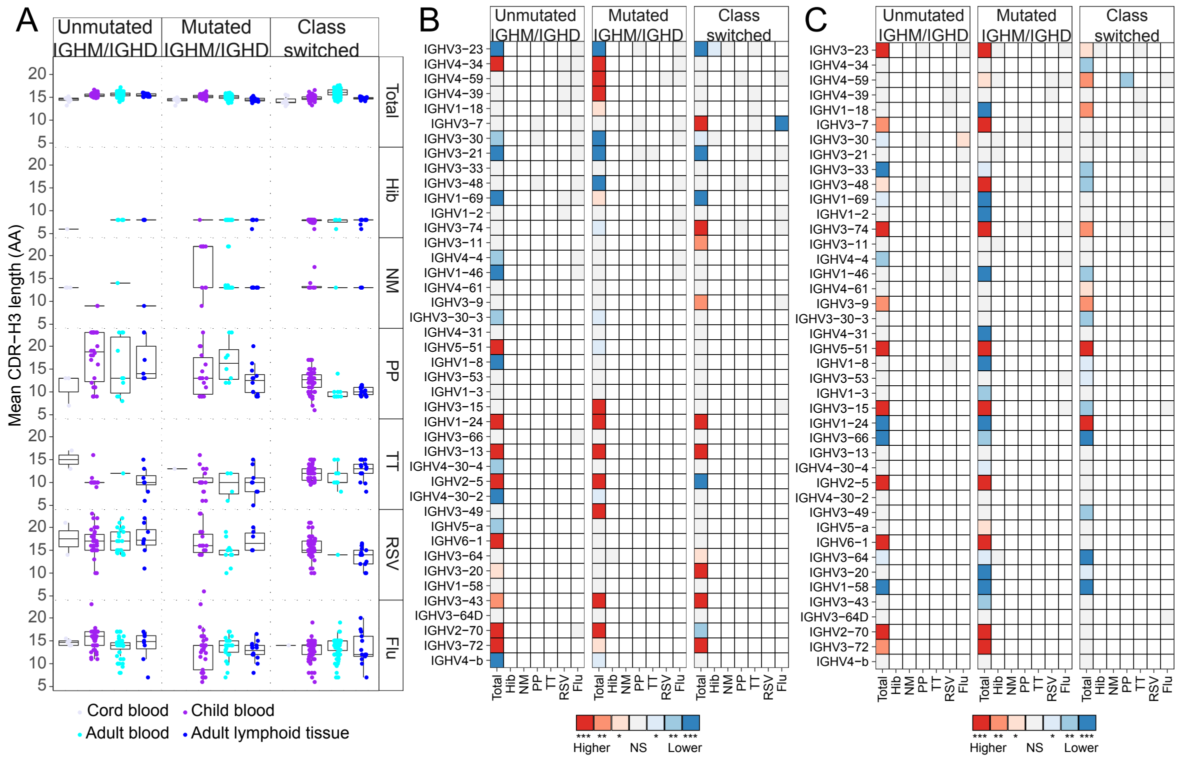


**Fig. S10. IGH CDR-H3 length and IGHV gene usage between samples of different age groups.** (A) The mean CDR-H3 length of total B cell clones (top panel) and convergent clones for each pathogen (row two to seven) in infant cord blood samples (lavender), children's blood (purple), adult blood (cyan) and adult lymphoid tissues (blue). Each dot represents the mean CDR-H3 length in a given sample. (B) Heatmap of IGHV gene usage of total B cells (left column) and convergent clones for each pathogen (labeled in columns) in children's blood compared to adult blood samples using paired Wilcoxon–Mann-Whitney test with Bonferroni correction. The color scale encodes the significance level and whether the usage was higher (red) or lower (blue) in children relative to adults. (C) Heatmap of IGHV gene usage of total B cells (left column) and convergent clones for each pathogen (right two to 7 columns) in adult blood compared to adult lymphoid tissue samples using paired Wilcoxon–Mann-Whitney test with Bonferroni correction. The color scale encodes the significance level and whether the usage was higher (red) or lower (blue) in lymphoid tissues relative to blood.

**Table S1. Demographic characteristics of deceased organ donors.**

| Donor ID | Sex | Age | Cause of death |
|---|---|---|---|
| Donor1 | M | 45 | Trauma |
| Donor2 | M | 25 | Anoxia |
| Donor3 | M | 31 | Trauma |
| Donor4 | M | 46 | Trauma |
| Donor5 | M | 32 | Anoxia, H1N1 2009 infection |
| Donor6 | F | 32 | Trauma |
| Donor7 | F | 62 | Cerebrovascular accident/ Stroke |
| Donor8 | F | 43 | Trauma |

All deceased organ donors were treated with steroids prior to organ recovery for transplantation.

**Table S2. References for public IGH sequences specific for each pathogen.**

| Pathogen | Source |
|---|---|
| Hib | 5. A. H. Lucas *et al.*, Molecular ontogeny of the human antibody repertoire to the Haemophilus influenzae type B polysaccharide: expression of canonical variable regions and their variants in vaccinated infants. *Clin Immunol* **108**, 119-127 (2003).<br>6. J. Truck *et al.*, Identification of antigen-specific B cell receptor sequences using public repertoire analysis. *J Immunol* **194**, 252-261 (2015).<br>7. E. E. Adderson *et al.*, Restricted immunoglobulin VH usage and VDJ combinations in the human response to Haemophilus influenzae type b capsular polysaccharide. Nucleotide sequences of monospecific anti-Haemophilus antibodies and polyspecific antibodies cross-reacting with self antigens. *J Clin Invest* **91**, 2734-2743 (1993).<br>8. A. H. Lucas, J. W. Larrick, D. C. Reason, Variable region sequences of a protective human monoclonal antibody specific for the Haemophilus influenzae type b capsular polysaccharide. *Infect Immun* **62**, 3873-3880 (1994). |
| NM | 9. J. D. Berry, D. J. Boese, D. K. Law, W. D. Zollinger, R. S. Tsang, Molecular analysis of monoclonal antibodies to group variant capsular polysaccharide of Neisseria meningitidis: recurrent heavy chains and alternative light chain partners. *Mol Immunol* **42**, 335-344 (2005).<br>10. S. L. Smithson, N. Srivastava, W. A. Hutchins, M. A. Westerink, Molecular analysis of the heavy chain of antibodies that recognize the capsular polysaccharide of Neisseria meningitidis in hu-PBMC reconstituted SCID mice and in the immunized human donor. *Mol Immunol* **36**, 113-124 (1999).<br>11. W. A. Hutchins, A. R. Adkins, T. Kieber-Emmons, M. A. Westerink, Molecular characterization of a monoclonal antibody produced in response to a group C meningococcal polysaccharide peptide mimic. *Mol Immunol* **33**, 503-510 (1996). |
| PP | 12. Z. Chen *et al.*, Human monoclonal antibodies isolated from a primary pneumococcal conjugate Vaccinee demonstrates the expansion of an antigen-driven Hypermutated memory B cell response. *BMC Infect Dis* **18**, 613 (2018).<br>13. K. Kolibab, S. L. Smithson, B. Rabquer, S. Khuder, M. A. Westerink, Immune response to pneumococcal polysaccharides 4 and 14 in elderly and young adults: analysis of the variable heavy chain repertoire. *Infect Immun* **73**, 7465-7476 (2005).<br>14. K. Smith *et al.*, Fully human monoclonal antibodies from antibody secreting cells after vaccination with Pneumovax(R)23 are serotype specific and facilitate opsonophagocytosis. *Immunobiology* **218**, 745-754 (2013). |

| | | |
|---|---|---|
| | 15. | A. S. Adler *et al.*, Rare, high-affinity anti-pathogen antibodies from human repertoires, discovered using microfluidics and molecular genomics. *MAbs* **9**, 1282-1296 (2017). |
| TT | 16. | B. J. DeKosky *et al.*, High-throughput sequencing of the paired human immunoglobulin heavy and light chain repertoire. *Nature biotechnology* **31**, 166-169 (2013). |
| | 17. | J. de Kruif *et al.*, Human immunoglobulin repertoires against tetanus toxoid contain a large and diverse fraction of high-affinity promiscuous V(H) genes. *J Mol Biol* **387**, 548-558 (2009). |
| | 18. | D. Frolich *et al.*, Secondary immunization generates clonally related antigen-specific plasma cells and memory B cells. *J Immunol* **185**, 3103-3110 (2010). |
| | 19. | T. R. Poulsen, A. Jensen, J. S. Haurum, P. S. Andersen, Limits for antibody affinity maturation and repertoire diversification in hypervaccinated humans. *J Immunol* **187**, 4229-4235 (2011). |
| RSV | 20. | M. S. Gilman *et al.*, Rapid profiling of RSV antibody repertoires from the memory B cells of naturally infected adult donors. *Sci Immunol* **1**, (2016). |
| | 21. | J. V. Williams, J. H. Weitkamp, D. L. Blum, B. J. LaFleur, J. E. Crowe, Jr., The human neonatal B cell response to respiratory syncytial virus uses a biased antibody variable gene repertoire that lacks somatic mutations. *Mol Immunol* **47**, 407-414 (2009). |
| | 22. | A. Tang *et al.*, A potent broadly neutralizing human RSV antibody targets conserved site IV of the fusion glycoprotein. *Nat Commun* **10**, 4153 (2019). |
| | 23. | B. Cortjens *et al.*, Broadly Reactive Anti-Respiratory Syncytial Virus G Antibodies from Exposed Individuals Effectively Inhibit Infection of Primary Airway Epithelial Cells. *J Virol* **91**, (2017). |
| | 24. | E. Goodwin *et al.*, Infants Infected with Respiratory Syncytial Virus Generate Potent Neutralizing Antibodies that Lack Somatic Hypermutation. *Immunity* **48**, 339-349 e335 (2018). |
| Flu | 25. | J. C. Krause *et al.*, Epitope-specific human influenza antibody repertoires diversify by B cell intraclonal sequence divergence and interclonal convergence. *J Immunol* **187**, 3704-3711 (2011). |
| | 26. | M. A. Moody *et al.*, H3N2 influenza infection elicits more cross-reactive and less clonally expanded anti-hemagglutinin antibodies than influenza vaccination. *PloS one* **6**, e25797 (2011). |
| | 27. | J. Wrammert *et al.*, Broadly cross-reactive antibodies dominate the human B cell response against 2009 pandemic H1N1 influenza virus infection. *J Exp Med* **208**, 181-193 (2011). |
| | 28. | A. H. Ellebedy *et al.*, Defining antigen-specific plasmablast and memory B cell subsets in human blood after viral infection or vaccination. *Nat Immunol* **17**, 1226-1234 (2016). |

29. D. Corti *et al.*, Tackling influenza with broadly neutralizing antibodies. *Curr Opin Virol* **24**, 60-69 (2017).
30. K. E. Neu *et al.*, Spec-seq unveils transcriptional subpopulations of antibody-secreting cells following influenza vaccination. *J Clin Invest* **129**, 93-105 (2019).
31. Y. Liu *et al.*, Cross-lineage protection by human antibodies binding the influenza B hemagglutinin. *Nat Commun* **10**, 324 (2019).
32. D. Hirano *et al.*, Three Types of Broadly Reacting Antibodies against Influenza B Viruses Induced by Vaccination with Seasonal Influenza Viruses. *J Immunol Res* **2018**, 7251793 (2018).
33. A. Yasuhara *et al.*, Diversity of antigenic mutants of influenza A(H1N1)pdm09 virus escaped from human monoclonal antibodies. *Scientific reports* **7**, 17735 (2017).
34. C. Shen *et al.*, A multimechanistic antibody targeting the receptor binding site potently cross-protects against influenza B viruses. *Sci Transl Med* **9**, (2017).
35. M. G. Joyce *et al.*, Vaccine-Induced Antibodies that Neutralize Group 1 and Group 2 Influenza A Viruses. *Cell* **166**, 609-623 (2016).
36. Y. Fu *et al.*, A broadly neutralizing anti-influenza antibody reveals ongoing capacity of haemagglutinin-specific memory B cells to evolve. *Nat Commun* **7**, 12780 (2016).
37. G. M. Li *et al.*, Pandemic H1N1 influenza vaccine induces a recall response in humans that favors broadly cross-reactive memory B cells. *Proceedings of the National Academy of Sciences of the United States of America* **109**, 9047-9052 (2012).
38. A. Watanabe *et al.*, Antibodies to a Conserved Influenza Head Interface Epitope Protect by an IgG Subtype-Dependent Mechanism. *Cell* **177**, 1124-1135 e1116 (2019).
39. K. R. McCarthy *et al.*, Memory B Cells that Cross-React with Group 1 and Group 2 Influenza A Viruses Are Abundant in Adult Human Repertoires. *Immunity* **48**, 174-184 e179 (2018).
40. S. F. Andrews *et al.*, Preferential induction of cross-group influenza A hemagglutinin stem-specific memory B cells after H7N9 immunization in humans. *Sci Immunol* **2**, (2017).

| | | |
|---|---|---|
| SARS-CoV-2 | 41. | P. J. M. Brouwer *et al.*, Potent neutralizing antibodies from COVID-19 patients define multiple targets of vulnerability. *Science*, (2020). |
| | 42. | A. Z. Wec *et al.*, Broad neutralization of SARS-related viruses by human monoclonal antibodies. *Science*, (2020). |
| | 43. | T. F. Rogers *et al.*, Isolation of potent SARS-CoV-2 neutralizing antibodies and protection from disease in a small animal model. *Science*, (2020). |
| | 44. | S. J. Zost *et al.*, Rapid isolation and profiling of a diverse panel of human monoclonal antibodies targeting the SARS-CoV-2 spike protein. *Nat Med*, (2020). |
| | 45. | R. Shi *et al.*, A human neutralizing antibody targets the receptor-binding site of SARS-CoV-2. *Nature*, (2020). |
| | 46. | E. Seydoux *et al.*, Characterization of neutralizing antibodies from a SARS-CoV-2 infected individual. *bioRxiv*, (2020). |
| | 47. | D. F. Robbiani *et al.*, Convergent Antibody Responses to SARS-CoV-2 Infection in Convalescent Individuals. *bioRxiv*, (2020). |
| | 48. | D. Pinto *et al.*, Structural and functional analysis of a potent sarbecovirus neutralizing antibody. *bioRxiv*, (2020). |
| | 49. | C. Kreer *et al.*, Longitudinal Isolation of Potent Near-Germline SARS-CoV-2-Neutralizing Antibodies from COVID-19 Patients. *Cell*, (2020). |
| EBOV | 50. | C. W. Davis *et al.*, Longitudinal Analysis of the Human B Cell Response to Ebola Virus Infection. *Cell* **177**, 1566-1582 e1517 (2019). |

**Table S3. Vaccine schedule for children in this study**

| ID | DTAP (Year_Month_Days) | HIB (Year_Month_Days) | PCV (Year_Month_Days) |
|---|---|---|---|
| 2800 | 0 Years 1 Months 11 Days | 0 Years 1 Months 11 Days | 0 Years 1 Months 11 Days |
| 2800 | 0 Years 3 Months 12 Days | 0 Years 3 Months 12 Days | 0 Years 3 Months 12 Days |
| 2800 | 0 Years 5 Months 30 Days | 0 Years 5 Months 30 Days | 0 Years 5 Months 30 Days |
| 2800 | 1 Years 3 Months 1 Days | 1 Years 3 Months 1 Days | 1 Years 3 Months 1 Days |
| 2801 | 0 Years 1 Months 13 Days | 0 Years 1 Months 13 Days | 0 Years 1 Months 13 Days |
| 2801 | 0 Years 3 Months 17 Days | 0 Years 3 Months 17 Days | 0 Years 3 Months 17 Days |
| 2801 | 0 Years 6 Months 2 Days | 0 Years 6 Months 2 Days | 0 Years 6 Months 2 Days |
| 2801 | 1 Years 9 Months 4 Days | 1 Years 9 Months 4 Days | 1 Years 9 Months 4 Days |
| 2802 | 0 Years 2 Months 2 Days | 0 Years 2 Months 2 Days | 0 Years 2 Months 2 Days |
| 2802 | 0 Years 4 Months 3 Days | 0 Years 4 Months 3 Days | 0 Years 4 Months 3 Days |
| 2802 | 0 Years 6 Months 13 Days | 0 Years 6 Months 13 Days | 0 Years 6 Months 13 Days |
| 2802 | 1 Years 3 Months 5 Days | | 1 Years 6 Months 6 Days |
| 2804 | 0 Years 1 Months 14 Days | 0 Years 1 Months 14 Days | 0 Years 1 Months 15 Days |
| 2804 | 0 Years 3 Months 18 Days | 0 Years 3 Months 18 Days | 0 Years 3 Months 18 Days |
| 2804 | 0 Years 6 Months 3 Days | 0 Years 6 Months 3 Days | 0 Years 6 Months 3 Days |
| 2804 | 1 Years 3 Months 15 Days | 1 Years 3 Months 15 Days | 1 Years 3 Months 15 Days |
| 2805 | 0 Years 2 Months 21 Days | 0 Years 2 Months 21 Days | 0 Years 2 Months 21 Days |
| 2805 | 0 Years 4 Months 2 Days | 0 Years 4 Months 2 Days | 0 Years 4 Months 2 Days |
| 2805 | 0 Years 7 Months 2 Days | 0 Years 7 Months 2 Days | 0 Years 7 Months 2 Days |
| 2805 | 1 Years 6 Months 13 Days | 1 Years 6 Months 13 Days | 1 Years 6 Months 13 Days |
| 2806 | 0 Years 1 Months 11 Days | 0 Years 1 Months 11 Days | 0 Years 1 Months 11 Days |
| 2806 | 0 Years 3 Months 18 Days | 0 Years 3 Months 18 Days | 0 Years 3 Months 18 Days |
| 2806 | 0 Years 7 Months 15 Days | 0 Years 7 Months 15 Days | 0 Years 7 Months 15 Days |
| 2806 | 1 Years 4 Months 30 Days | 1 Years 4 Months 30 Days | 1 Years 4 Months 30 Days |
| 2807 | 0 Years 2 Months 1 Days | 0 Years 2 Months 1 Days | 0 Years 2 Months 1 Days |
| 2807 | 0 Years 4 Months 2 Days | 0 Years 4 Months 2 Days | 0 Years 4 Months 2 Days |
| 2807 | 0 Years 6 Months 3 Days | 0 Years 6 Months 3 Days | 0 Years 6 Months 3 Days |
| 2807 | 1 Years 2 Months 5 Days | 1 Years 2 Months 5 Days | 1 Years 2 Months 5 Days |
| 2808 | 0 Years 1 Months 23 Days | 0 Years 1 Months 23 Days | 0 Years 2 Months 4 Days |
| 2808 | 0 Years 4 Months 9 Days | 0 Years 4 Months 9 Days | 0 Years 4 Months 9 Days |
| 2808 | 0 Years 6 Months 11 Days | 0 Years 6 Months 11 Days | 0 Years 6 Months 11 Days |
| 2808 | 1 Years 0 Months 24 Days | 1 Years 3 Months 28 Days | 1 Years 3 Months 28 Days |
| 2809 | 0 Years 2 Months 10 Days | 0 Years 2 Months 10 Days | 0 Years 2 Months 10 Days |
| 2809 | 0 Years 4 Months 13 Days | 0 Years 4 Months 13 Days | 0 Years 4 Months 13 Days |
| 2809 | 0 Years 6 Months 8 Days | 0 Years 6 Months 8 Days | 0 Years 6 Months 8 Days |
| 2809 | 1 Years 3 Months 17 Days | 1 Years 3 Months 17 Days | 1 Years 3 Months 17 Days |
| 2810 | | 1 Years 0 Months 3 Days | 1 Years 0 Months 3 Days |

| | | | |
|---|---|---|---|
| 2811 | 0 Years 3 Months 6 Days | 0 Years 3 Months 6 Days | 0 Years 3 Months 6 Days |
| 2811 | 0 Years 5 Months 14 Days | 0 Years 5 Months 14 Days | 0 Years 5 Months 14 Days |
| 2811 | 0 Years 7 Months 14 Days | 0 Years 7 Months 14 Days | 0 Years 7 Months 14 Days |
| 2811 | 1 Years 3 Months 12 Days | 1 Years 0 Months 4 Days | 1 Years 0 Months 4 Days |
| 2812 | 0 Years 2 Months 11 Days | 0 Years 2 Months 11 Days | 0 Years 2 Months 11 Days |
| 2812 | 0 Years 5 Months 3 Days | 0 Years 5 Months 3 Days | 0 Years 5 Months 3 Days |
| 2812 | 0 Years 7 Months 5 Days | 0 Years 7 Months 5 Days | 0 Years 7 Months 5 Days |
| 2812 | 1 Years 4 Months 16 Days | 1 Years 4 Months 16 Days | 1 Years 1 Months 14 Days |
| 2813 | 0 Years 1 Months 18 Days | 0 Years 1 Months 18 Days | 0 Years 1 Months 18 Days |
| 2813 | 0 Years 6 Months 1 Days | 0 Years 6 Months 1 Days | 0 Years 6 Months 1 Days |
| 2813 | 0 Years 8 Months 6 Days | 0 Years 8 Months 6 Days | 1 Years 0 Months 13 Days |
| 2813 | 1 Years 3 Months 18 Days | 1 Years 3 Months 18 Days | |
| 2814 | 0 Years 1 Months 27 Days | 0 Years 1 Months 27 Days | 0 Years 1 Months 27 Days |
| 2814 | 0 Years 3 Months 30 Days | 0 Years 3 Months 30 Days | 0 Years 3 Months 30 Days |
| 2814 | 0 Years 6 Months 19 Days | 0 Years 6 Months 19 Days | 0 Years 6 Months 19 Days |
| 2814 | 2 Years 4 Months 16 Days | 2 Years 4 Months 16 Days | 3 Years 0 Months 29 Days |
| 2815 | 0 Years 3 Months 30 Days | 0 Years 3 Months 30 Days | 0 Years 3 Months 30 Days |
| 2815 | 0 Years 6 Months 30 Days | 0 Years 6 Months 30 Days | 0 Years 6 Months 30 Days |
| 2815 | 1 Years 5 Months 21 Days | | |
| 2816 | 0 Years 2 Months 3 Days | 0 Years 2 Months 3 Days | 0 Years 2 Months 3 Days |
| 2816 | 0 Years 4 Months 25 Days | 0 Years 4 Months 25 Days | 0 Years 4 Months 25 Days |
| 2816 | 0 Years 6 Months 27 Days | 0 Years 6 Months 27 Days | 0 Years 6 Months 27 Days |
| 2816 | 1 Years 4 Months 18 Days | 1 Years 4 Months 18 Days | 1 Years 0 Months 1 Days |
| 2817 | 0 Years 2 Months 4 Days | 0 Years 2 Months 4 Days | 0 Years 2 Months 4 Days |
| 2817 | 0 Years 6 Months 1 Days | 0 Years 6 Months 1 Days | 0 Years 4 Months 2 Days |
| 2817 | 0 Years 6 Months 1 Days | 0 Years 6 Months 1 Days | 0 Years 6 Months 1 Days |
| 2817 | 1 Years 3 Months 1 Days | 1 Years 0 Months 6 Days | 1 Years 0 Months 6 Days |
| 2819 | 0 Years 2 Months 3 Days | 0 Years 2 Months 3 Days | 0 Years 2 Months 3 Days |
| 2819 | 0 Years 4 Months 2 Days | 0 Years 4 Months 2 Days | 0 Years 4 Months 2 Days |
| 2819 | 0 Years 5 Months 27 Days | 0 Years 5 Months 27 Days | 0 Years 5 Months 27 Days |
| 2819 | 1 Years 3 Months 20 Days | 1 Years 0 Months 17 Days | 1 Years 0 Months 17 Days |
| 2820 | 0 Years 0 Months 24 Days | 0 Years 0 Months 24 Days | 0 Years 0 Months 24 Days |
| 2820 | 0 Years 2 Months 7 Days | 0 Years 2 Months 7 Days | 0 Years 2 Months 7 Days |
| 2821 | 0 Years 5 Months 23 Days | 0 Years 5 Months 23 Days | 0 Years 5 Months 23 Days |
| 2821 | 0 Years 7 Months 23 Days | 0 Years 7 Months 23 Days | 0 Years 7 Months 23 Days |
| 2821 | 0 Years 11 Months 7 Days | 0 Years 10 Months 24 Days | 0 Years 10 Months 24 Days |
| 2821 | 1 Years 4 Months 30 Days | 1 Years 4 Months 30 Days | 1 Years 1 Months 26 Days |
| 2822 | 0 Years 3 Months 29 Days | 0 Years 3 Months 29 Days | 0 Years 3 Months 29 Days |
| 2822 | 0 Years 8 Months 16 Days | 0 Years 8 Months 16 Days | 0 Years 8 Months 16 Days |

| | | | |
|---|---|---|---|
| 2822 | 1 Years 0 Months 1 Days | 1 Years 0 Months 1 Days | 1 Years 0 Months 1 Days |
| 2822 | 1 Years 5 Months 18 Days | 1 Years 5 Months 18 Days | 1 Years 5 Months 18 Days |
| 3550 | 0 Years 1 Months 25 Days | 0 Years 1 Months 25 Days | 0 Years 1 Months 25 Days |
| 3550 | 0 Years 3 Months 27 Days | 0 Years 3 Months 27 Days | 0 Years 3 Months 27 Days |
| 3550 | 0 Years 6 Months 3 Days | 0 Years 6 Months 3 Days | 0 Years 6 Months 3 Days |
| 3551 | 0 Years 1 Months 11 Days | 0 Years 1 Months 11 Days | 0 Years 1 Months 11 Days |
| 3551 | 0 Years 3 Months 6 Days | 0 Years 3 Months 6 Days | 0 Years 3 Months 6 Days |
| 3551 | 0 Years 6 Months 1 Days | 0 Years 6 Months 1 Days | 0 Years 6 Months 1 Days |
| 3551 | 1 Years 4 Months 26 Days | 1 Years 4 Months 26 Days | 1 Years 4 Months 26 Days |
| 3552 | 0 Years 1 Months 20 Days | 0 Years 1 Months 20 Days | 0 Years 1 Months 20 Days |
| 3552 | 0 Years 6 Months 6 Days | 0 Years 6 Months 6 Days | 0 Years 6 Months 6 Days |
| 3552 | 1 Years 3 Months 21 Days | 1 Years 3 Months 21 Days | 1 Years 3 Months 21 Days |
| 3553 | 0 Years 2 Months 9 Days | 0 Years 2 Months 9 Days | 0 Years 2 Months 9 Days |
| 3553 | 0 Years 4 Months 4 Days | 0 Years 4 Months 4 Days | 0 Years 4 Months 4 Days |
| 3553 | 0 Years 6 Months 3 Days | 0 Years 6 Months 3 Days | 0 Years 6 Months 3 Days |
| 3553 | 1 Years 3 Months 5 Days | 1 Years 0 Months 8 Days | 1 Years 0 Months 8 Days |
| 3554 | 0 Years 1 Months 11 Days | 0 Years 1 Months 11 Days | 0 Years 1 Months 11 Days |
| 3554 | 0 Years 5 Months 1 Days | 0 Years 5 Months 1 Days | 0 Years 5 Months 1 Days |
| 3554 | 0 Years 7 Months 3 Days | 0 Years 7 Months 3 Days | 0 Years 7 Months 3 Days |
| 3554 | 1 Years 2 Months 29 Days | 1 Years 2 Months 29 Days | 1 Years 2 Months 29 Days |
| 3556 | 0 Years 1 Months 13 Days | 0 Years 1 Months 13 Days | 0 Years 1 Months 13 Days |
| 3556 | 0 Years 3 Months 12 Days | 0 Years 3 Months 12 Days | 0 Years 3 Months 12 Days |
| 3556 | 0 Years 5 Months 29 Days | 0 Years 5 Months 29 Days | 0 Years 5 Months 29 Days |
| 3556 | 1 Years 7 Months 15 Days | 1 Years 7 Months 15 Days | 1 Years 7 Months 15 Days |
| 3557 | 0 Years 1 Months 29 Days | 0 Years 1 Months 29 Days | 0 Years 1 Months 29 Days |
| 3558 | 0 Years 2 Months 3 Days | 0 Years 2 Months 3 Days | 0 Years 2 Months 3 Days |
| 3558 | 0 Years 5 Months 2 Days | 0 Years 5 Months 2 Days | 0 Years 5 Months 2 Days |
| 3558 | 0 Years 6 Months 19 Days | 0 Years 6 Months 19 Days | 0 Years 6 Months 19 Days |
| 3558 | 1 Years 3 Months 6 Days | 1 Years 0 Months 3 Days | |
| 3559 | 4 Years 1 Months 5 Days | | |
| 3559 | 4 Years 7 Months 29 Days | | |
| 3560 | 0 Years 1 Months 16 Days | 0 Years 1 Months 16 Days | 0 Years 1 Months 16 Days |
| 3560 | 0 Years 3 Months 17 Days | 0 Years 3 Months 17 Days | 0 Years 3 Months 17 Days |
| 3560 | 0 Years 6 Months 6 Days | 0 Years 6 Months 6 Days | 0 Years 6 Months 6 Days |
| 3560 | 1 Years 2 Months 29 Days | 1 Years 2 Months 29 Days | 1 Years 2 Months 29 Days |
| 3561 | 0 Years 2 Months 1 Days | 0 Years 2 Months 1 Days | 0 Years 2 Months 1 Days |
| 3561 | 0 Years 4 Months 15 Days | 0 Years 4 Months 15 Days | 0 Years 4 Months 15 Days |
| 3561 | 0 Years 5 Months 22 Days | 0 Years 5 Months 22 Days | 0 Years 5 Months 22 Days |
| 3561 | 1 Years 3 Months 18 Days | | 1 Years 2 Months 14 Days |

| | | | |
|---|---|---|---|
| 3563 | 0 Years 1 Months 29 Days | 0 Years 1 Months 29 Days | 0 Years 1 Months 29 Days |
| 3563 | 0 Years 4 Months 12 Days | 0 Years 4 Months 12 Days | 0 Years 4 Months 12 Days |
| 3563 | 0 Years 6 Months 26 Days | 0 Years 6 Months 26 Days | 0 Years 6 Months 26 Days |
| 3563 | 1 Years 3 Months 25 Days | 1 Years 0 Months 5 Days | 1 Years 0 Months 5 Days |
| 3564 | 0 Years 2 Months 5 Days | 0 Years 2 Months 5 Days | 0 Years 2 Months 5 Days |
| 3564 | 0 Years 4 Months 11 Days | 0 Years 4 Months 11 Days | 0 Years 4 Months 11 Days |
| 3564 | 0 Years 6 Months 14 Days | 0 Years 6 Months 14 Days | 0 Years 6 Months 14 Days |
| 3564 | 1 Years 3 Months 15 Days | 1 Years 0 Months 5 Days | 1 Years 0 Months 5 Days |
| 3565 | 0 Years 2 Months 1 Days | 0 Years 2 Months 1 Days | 0 Years 2 Months 1 Days |
| 3565 | 0 Years 4 Months 3 Days | 0 Years 4 Months 3 Days | 0 Years 4 Months 3 Days |
| 3565 | 0 Years 5 Months 27 Days | 0 Years 5 Months 27 Days | 0 Years 5 Months 27 Days |
| 3565 | 1 Years 4 Months 15 Days | 1 Years 4 Months 15 Days | 1 Years 4 Months 15 Days |
| 3960 | 0 Years 2 Months 4 Days | 0 Years 2 Months 4 Days | 0 Years 3 Months 30 Days |
| 3960 | 0 Years 3 Months 30 Days | 0 Years 3 Months 30 Days | 0 Years 2 Months 4 Days |
| 3960 | 0 Years 6 Months 2 Days | 0 Years 6 Months 2 Days | 0 Years 6 Months 2 Days |
| 3960 | 1 Years 0 Months 2 Days | 1 Years 0 Months 2 Days | 1 Years 0 Months 2 Days |
| 3961 | 0 Years 1 Months 29 Days | 0 Years 1 Months 29 Days | 0 Years 1 Months 29 Days |
| 3961 | 0 Years 4 Months 2 Days | 0 Years 4 Months 2 Days | 0 Years 4 Months 2 Days |
| 3961 | 0 Years 6 Months 28 Days | 0 Years 6 Months 28 Days | 0 Years 6 Months 28 Days |
| 3961 | 1 Years 3 Months 14 Days | 1 Years 2 Months 6 Days | 1 Years 2 Months 6 Days |
| 3962 | 0 Years 2 Months 8 Days | 0 Years 2 Months 8 Days | 0 Years 2 Months 8 Days |
| 3962 | 0 Years 3 Months 29 Days | 0 Years 3 Months 29 Days | 0 Years 3 Months 29 Days |
| 3962 | 0 Years 6 Months 2 Days | 0 Years 6 Months 2 Days | 0 Years 6 Months 2 Days |
| 3962 | 1 Years 2 Months 24 Days | 1 Years 2 Months 24 Days | 1 Years 0 Months 3 Days |
| 3963 | 0 Years 1 Months 20 Days | 0 Years 1 Months 20 Days | 0 Years 1 Months 20 Days |
| 3963 | 0 Years 3 Months 21 Days | 0 Years 3 Months 21 Days | 0 Years 3 Months 21 Days |
| 3963 | 0 Years 6 Months 29 Days | 0 Years 6 Months 29 Days | 0 Years 6 Months 29 Days |
| 3963 | 1 Years 3 Months 2 Days | 1 Years 3 Months 2 Days | 1 Years 3 Months 2 Days |
| 3964 | 0 Years 1 Months 20 Days | 0 Years 1 Months 20 Days | 0 Years 1 Months 20 Days |
| 3964 | 0 Years 3 Months 23 Days | 0 Years 3 Months 23 Days | 0 Years 3 Months 23 Days |
| 3964 | 0 Years 6 Months 9 Days | 0 Years 6 Months 9 Days | 0 Years 6 Months 9 Days |
| 3964 | 1 Years 4 Months 29 Days | 1 Years 4 Months 29 Days | 1 Years 4 Months 29 Days |
| 3965 | 0 Years 1 Months 11 Days | 0 Years 1 Months 11 Days | 0 Years 1 Months 11 Days |
| 3965 | 0 Years 3 Months 13 Days | 0 Years 3 Months 13 Days | 0 Years 3 Months 13 Days |
| 3965 | 0 Years 6 Months 21 Days | 0 Years 6 Months 21 Days | 0 Years 6 Months 21 Days |
| 3965 | 1 Years 4 Months 12 Days | 1 Years 4 Months 12 Days | 1 Years 4 Months 12 Days |
| 3966 | 0 Years 2 Months 8 Days | 0 Years 2 Months 8 Days | 0 Years 2 Months 8 Days |
| 3966 | 0 Years 4 Months 10 Days | 0 Years 4 Months 10 Days | 0 Years 4 Months 10 Days |
| 3966 | 0 Years 6 Months 15 Days | 0 Years 6 Months 15 Days | 0 Years 9 Months 5 Days |

| | | | |
|---|---|---|---|
| 3966 | 1 Years 3 Months 1 Days | 1 Years 3 Months 1 Days | 1 Years 3 Months 1 Days |
| 3967 | 0 Years 2 Months 3 Days | 0 Years 2 Months 3 Days | 0 Years 2 Months 3 Days |
| 3967 | 0 Years 4 Months 4 Days | 0 Years 4 Months 4 Days | 0 Years 4 Months 4 Days |
| 3967 | 0 Years 6 Months 2 Days | 0 Years 6 Months 2 Days | 0 Years 6 Months 2 Days |
| 3967 | 1 Years 3 Months 12 Days | 1 Years 4 Months 13 Days | 1 Years 6 Months 5 Days |
| 4680 | 0 Years 2 Months 6 Days | 0 Years 2 Months 6 Days | 0 Years 2 Months 6 Days |
| 4680 | 0 Years 4 Months 1 Days | 0 Years 4 Months 1 Days | 0 Years 4 Months 1 Days |
| 4680 | 0 Years 5 Months 26 Days | 0 Years 5 Months 26 Days | 0 Years 5 Months 26 Days |
| 4680 | 1 Years 3 Months 7 Days | 1 Years 0 Months 1 Days | 1 Years 0 Months 1 Days |
| 4681 | 0 Years 1 Months 27 Days | 0 Years 1 Months 27 Days | 0 Years 1 Months 27 Days |
| 4681 | 0 Years 3 Months 30 Days | 0 Years 3 Months 30 Days | 0 Years 3 Months 30 Days |
| 4681 | 0 Years 6 Months 2 Days | 0 Years 6 Months 2 Days | 0 Years 6 Months 2 Days |
| 4682 | 0 Years 1 Months 22 Days | 0 Years 1 Months 22 Days | 0 Years 1 Months 22 Days |
| 4682 | 0 Years 5 Months 21 Days | 0 Years 5 Months 21 Days | 0 Years 5 Months 21 Days |
| 4682 | 0 Years 7 Months 16 Days | 0 Years 7 Months 16 Days | 0 Years 7 Months 16 Days |
| 4682 | 1 Years 6 Months 16 Days | 1 Years 6 Months 16 Days | 1 Years 0 Months 4 Days |
| 4683 | 0 Years 1 Months 27 Days | 0 Years 1 Months 27 Days | 0 Years 1 Months 27 Days |
| 4683 | 0 Years 3 Months 28 Days | 0 Years 3 Months 28 Days | 0 Years 3 Months 28 Days |
| 4683 | 0 Years 6 Months 3 Days | 0 Years 6 Months 3 Days | 0 Years 6 Months 3 Days |
| 4683 | 1 Years 3 Months 5 Days | 1 Years 3 Months 5 Days | 1 Years 3 Months 5 Days |

DTAP: diphtheria, tetanus, and acellular pertussis vaccine
HIB: Haemophilus influenzae type B vaccine
PCV: Pneumococcal vaccine

**Table S4. List of reported SARS-CoV-2 antibodies that can be found in child blood samples and which can bind to the RBD of SARS-CoV-2, except neutralizing antibodies and RBD-binding antibodies which are listed in Table 1**

| ID | Vgene | Jgene | CDR-H3 | Protein; Epitope |
|---|---|---|---|---|
| COVA2-14 | IGHV1-69 | IGHJ4 | ARVRYYDSSGYYEDY | S; non-RBD |
| COVA3-01 | IGHV4-59 | IGHJ6 | ARGPAATYYYYMDV | S; non-RBD |
| CC12.25 | IGHV3-23 | IGHJ5 | AKDRYYEFWSGYSNWFDP | S; non-RBD |
| COV2-2011 | IGHV3-30-3 | IGHJ6 | ARGHTGNYYYGMDV | S; non-RBD |
| COV2-2151 | IGHV1-69 | IGHJ1 | ARIGSYPEYFQH | S; non-RBD |
| COV2-2159 | IGHV3-30-3 | IGHJ6 | ARSTSGSYYYGMDV | S; non-RBD |
| COV2-2160 | IGHV3-30-3 | IGHJ6 | ARSTSGSYYYGMDV | S; non-RBD |
| COV2-2178 | IGHV3-7 | IGHJ4 | ARVGSSSWYFDY | S; non-RBD |
| COV2-2341 | IGHV3-30-3 | IGHJ6 | ARSTSGSYYYGMDV | S; non-RBD |
| COV2-2517 | IGHV1-69 | IGHJ1 | ARIGSYPEYFQH | S; non-RBD |
| COV2-2564 | IGHV3-30-3 | IGHJ6 | ARAQGGNYYYGMDV | S; non-RBD |
| COV2-2844 | IGHV3-30-3 | IGHJ6 | ARAQGGNYYYGMDV | S; non-RBD |
| COVA1-02 | IGHV3-30 | IGHJ3 | ARARGGSYNDAFDI | S; non-RBD |
| COVA2-34 | IGHV3-30 | IGHJ4 | ARSASGSYYGAFDY | S; non-RBD |
| COV2-2143 | IGHV3-66 | IGHJ6 | AKEGGSGSLRYYYYGMDV | S; non-RBD |
| COV2-2147 | IGHV3-30-3 | IGHJ6 | ARSTSGSYYYGMDV | S; non-RBD |
| COV2-2386 | IGHV3-33 | IGHJ5 | AREGDFWSGYYTGWFDP | S; non-RBD |
| COV2-2418 | IGHV3-23 | IGHJ6 | AKPYGMDV | S; non-RBD |
| COV2-2675 | IGHV3-30 | IGHJ5 | AKDGSGSYYGWFDP | S; non-RBD |
| CV10 | IGHV4-59 | IGHJ4 | ARGFDY | S; non-RBD |
| CV2 | IGHV3-30 | IGHJ4 | ARVRGSYYLFDY | S; non-RBD |
| CV25 | IGHV4-30-4 | IGHJ6 | ARDHHYDFWSGYSSYYYYGMDV | S; non-RBD |
| CV27 | IGHV3-30 | IGHJ6 | ARSFGGSYYYGMDV | S; non-RBD |
| CV33 | IGHV1-18 | IGHJ6 | ARDSVAGIYYYYGMDV | S; non-RBD |
| CV34 | IGHV3-30-3 | IGHJ6 | ARSYGGSYYYGMDV | S; non-RBD |
| COV2-2622 | IGHV4-4 | IGHJ4 | ARGWYFDY | S; NTD |
| mAb-162 | IGHV3-30 | IGHJ4 | ARDLPPLDY | S; Unk |
| mAb-154 | IGHV1-2 | IGHJ4 | ASGPNYFDY | S; Unk |
| mAb-42 | IGHV3-30 | IGHJ4 | ARDLPPLDY | S; Unk |
| CC12.24 | IGHV3-30 | IGHJ6 | AKDRTGNYYYGMDV | S; Unk |

**Table S5. Primer sequences**

| FR1 primer sets | Primer sequences |
|---|---|
| P7_VH1_FR1_A1 | GGCATTCCTGCTGAACCGCTCTTCCGATCTatgatacaGGCCTCAGTGAAGGTCTCCTGCAAG |
| P7_VH2_FR1_A1 | GGCATTCCTGCTGAACCGCTCTTCCGATCTatgatacaGTCTGGTCCTACGCTGGTGAAACCC |
| P7_VH3_FR1_A1 | GGCATTCCTGCTGAACCGCTCTTCCGATCTatgatacaCTGGGGGGTCCCTGAGACTCTCCTG |
| P7_VH4_FR1_A1 | GGCATTCCTGCTGAACCGCTCTTCCGATCTatgatacaCTTCGGAGACCCTGTCCCTCACCTG |
| P7_VH5_FR1_A1 | GGCATTCCTGCTGAACCGCTCTTCCGATCTatgatacaCGGGGAGTCTCTGAAGATCTCCTGT |
| P7_VH6_FR1_A1 | GGCATTCCTGCTGAACCGCTCTTCCGATCTatgatacaTCGCAGACCCTCTCACTCACCTGTG |
| P7_VH1_FR1_A2 | GGCATTCCTGCTGAACCGCTCTTCCGATCTcatgtcatGGCCTCAGTGAAGGTCTCCTGCAAG |
| P7_VH2_FR1_A2 | GGCATTCCTGCTGAACCGCTCTTCCGATCTcatgtcatGTCTGGTCCTACGCTGGTGAAACCC |
| P7_VH3_FR1_A2 | GGCATTCCTGCTGAACCGCTCTTCCGATCTcatgtcatCTGGGGGGTCCCTGAGACTCTCCTG |
| P7_VH4_FR1_A2 | GGCATTCCTGCTGAACCGCTCTTCCGATCTcatgtcatCTTCGGAGACCCTGTCCCTCACCTG |
| P7_VH5_FR1_A2 | GGCATTCCTGCTGAACCGCTCTTCCGATCTcatgtcatCGGGGAGTCTCTGAAGATCTCCTGT |
| P7_VH6_FR1_A2 | GGCATTCCTGCTGAACCGCTCTTCCGATCTcatgtcatTCGCAGACCCTCTCACTCACCTGTG |
| P7_VH1_FR1_A3 | GGCATTCCTGCTGAACCGCTCTTCCGATCTcacgcactGGCCTCAGTGAAGGTCTCCTGCAAG |
| P7_VH2_FR1_A3 | GGCATTCCTGCTGAACCGCTCTTCCGATCTcacgcactGTCTGGTCCTACGCTGGTGAAACCC |
| P7_VH3_FR1_A3 | GGCATTCCTGCTGAACCGCTCTTCCGATCTcacgcactCTGGGGGGTCCCTGAGACTCTCCTG |
| P7_VH4_FR1_A3 | GGCATTCCTGCTGAACCGCTCTTCCGATCTcacgcactCTTCGGAGACCCTGTCCCTCACCTG |
| P7_VH5_FR1_A3 | GGCATTCCTGCTGAACCGCTCTTCCGATCTcacgcactCGGGGAGTCTCTGAAGATCTCCTGT |
| P7_VH6_FR1_A3 | GGCATTCCTGCTGAACCGCTCTTCCGATCTcacgcactTCGCAGACCCTCTCACTCACCTGTG |
| P7_VH1_FR1_A4 | GGCATTCCTGCTGAACCGCTCTTCCGATCTtgtgagcgGGCCTCAGTGAAGGTCTCCTGCAAG |
| P7_VH2_FR1_A4 | GGCATTCCTGCTGAACCGCTCTTCCGATCTtgtgagcgGTCTGGTCCTACGCTGGTGAAACCC |
| P7_VH3_FR1_A4 | GGCATTCCTGCTGAACCGCTCTTCCGATCTtgtgagcgCTGGGGGGTCCCTGAGACTCTCCTG |

| | |
|---|---|
| P7_VH4_FR1_A4 | GGCATTCCTGCTGAACCGCTCTTCCGATCTtgtgagcgCTTCGGAG ACCCTGTCCCTCACCTG |
| P7_VH5_FR1_A4 | GGCATTCCTGCTGAACCGCTCTTCCGATCTtgtgagcgCGGGGAGT CTCTGAAGATCTCCTGT |
| P7_VH6_FR1_A4 | GGCATTCCTGCTGAACCGCTCTTCCGATCTtgtgagcgTCGCAGAC CCTCTCACTCACCTGTG |
| Isotype primer | Primer sequences |
| P5_IgG_long_A1 | ACACTCTTTCCCTACACGACGCTCTTCCGATCTNNNNATGATA CACAGGCAGCCCAGGGC |
| P5_IgA_long_A1 | ACACTCTTTCCCTACACGACGCTCTTCCGATCTNNNNATGATA CAAGCCCTGGACCAGGCA |
| P5_IgD_A1 | ACACTCTTTCCCTACACGACGCTCTTCCGATCTNNNNATGATA CACCCTGATATGATGGGGAACA |
| P5_IgM_A1 | ACACTCTTTCCCTACACGACGCTCTTCCGATCTNNNNATGATA CAGGGAATTCTCACAGGAGACG |
| P5_IgE_A1 | ACACTCTTTCCCTACACGACGCTCTTCCGATCTNNNNATGATA CAGAAGACGGATGGGCTCTGT |
| P5_IgG_long_A2 | ACACTCTTTCCCTACACGACGCTCTTCCGATCTNNNNCATGTC ATCAGGCAGCCCAGGGC |
| P5_IgA_long_A2 | ACACTCTTTCCCTACACGACGCTCTTCCGATCTNNNNCATGTC ATAGCCCTGGACCAGGCA |
| P5_IgD_A2 | ACACTCTTTCCCTACACGACGCTCTTCCGATCTNNNNCATGTC ATCCCTGATATGATGGGGAACA |
| P5_IgM_A2 | ACACTCTTTCCCTACACGACGCTCTTCCGATCTNNNNCATGTC ATGGGAATTCTCACAGGAGACG |
| P5_IgE_A2 | ACACTCTTTCCCTACACGACGCTCTTCCGATCTNNNNCATGTC ATGAAGACGGATGGGCTCTGT |
| P5_IgG_long_A3 | ACACTCTTTCCCTACACGACGCTCTTCCGATCTNNNNCACGCA CTCAGGCAGCCCAGGGC |
| P5_IgA_long_A3 | ACACTCTTTCCCTACACGACGCTCTTCCGATCTNNNNCACGCA CTAGCCCTGGACCAGGCA |
| P5_IgD_A3 | ACACTCTTTCCCTACACGACGCTCTTCCGATCTNNNNCACGCA CTCCCTGATATGATGGGGAACA |
| P5_IgM_A3 | ACACTCTTTCCCTACACGACGCTCTTCCGATCTNNNNCACGCA CTGGGAATTCTCACAGGAGACG |
| P5_IgE_A3 | ACACTCTTTCCCTACACGACGCTCTTCCGATCTNNNNCACGCA CTGAAGACGGATGGGCTCTGT |
| P5_IgG_long_A4 | ACACTCTTTCCCTACACGACGCTCTTCCGATCTNNNNTGTGAG CGCAGGCAGCCCAGGGC |
| P5_IgA_long_A4 | ACACTCTTTCCCTACACGACGCTCTTCCGATCTNNNNTGTGAG CGAGCCCTGGACCAGGCA |
| P5_IgD_A4 | ACACTCTTTCCCTACACGACGCTCTTCCGATCTNNNNTGTGAG CGCCCTGATATGATGGGGAACA |
| P5_IgM_A4 | ACACTCTTTCCCTACACGACGCTCTTCCGATCTNNNNTGTGAG CGGGGAATTCTCACAGGAGACG |

| | |
|---|---|
| P5_IgE_A4 | ACACTCTTTCCCTACACGACGCTCTTCCGATCTNNNNTGTGAGCGGAAGACGGATGGGCTCTGT |
| P5_IgG_long_A5 | ACACTCTTTCCCTACACGACGCTCTTCCGATCTNNNNGCGCATGCCAGGCAGCCCAGGGC |
| P5_IgA_long_A5 | ACACTCTTTCCCTACACGACGCTCTTCCGATCTNNNNGCGCATGCAGCCCTGGACCAGGCA |
| P5_IgD_A5 | ACACTCTTTCCCTACACGACGCTCTTCCGATCTNNNNGCGCATGCCCCTGATATGATGGGGAACA |
| P5_IgM_A5 | ACACTCTTTCCCTACACGACGCTCTTCCGATCTNNNNGCGCATGCGGGAATTCTCACAGGAGACG |
| P5_IgE_A5 | ACACTCTTTCCCTACACGACGCTCTTCCGATCTNNNNGCGCATGCGAAGACGGATGGGCTCTGT |
| P5_IgG_long_A6 | ACACTCTTTCCCTACACGACGCTCTTCCGATCTNNNNACACGACGCAGGCAGCCCAGGGC |
| P5_IgA_long_A6 | ACACTCTTTCCCTACACGACGCTCTTCCGATCTNNNNACACGACGAGCCCTGGACCAGGCA |
| P5_IgD_A6 | ACACTCTTTCCCTACACGACGCTCTTCCGATCTNNNNACACGACGCCCTGATATGATGGGGAACA |
| P5_IgM_A6 | ACACTCTTTCCCTACACGACGCTCTTCCGATCTNNNNACACGACGGGGAATTCTCACAGGAGACG |
| P5_IgE_A6 | ACACTCTTTCCCTACACGACGCTCTTCCGATCTNNNNACACGACGGAAGACGGATGGGCTCTGT |
| P5_IgG_long_A7 | ACACTCTTTCCCTACACGACGCTCTTCCGATCTNNNNTCGTCGTCCAGGCAGCCCAGGGC |
| P5_IgA_long_A7 | ACACTCTTTCCCTACACGACGCTCTTCCGATCTNNNNTCGTCGTCAGCCCTGGACCAGGCA |
| P5_IgD_A7 | ACACTCTTTCCCTACACGACGCTCTTCCGATCTNNNNTCGTCGTCCCCTGATATGATGGGGAACA |
| P5_IgM_A7 | ACACTCTTTCCCTACACGACGCTCTTCCGATCTNNNNTCGTCGTCGGGAATTCTCACAGGAGACG |
| P5_IgE_A7 | ACACTCTTTCCCTACACGACGCTCTTCCGATCTNNNNTCGTCGTCGAAGACGGATGGGCTCTGT |
| P5_IgG_long_A8 | ACACTCTTTCCCTACACGACGCTCTTCCGATCTNNNNCGTATGCACAGGCAGCCCAGGGC |
| P5_IgA_long_A8 | ACACTCTTTCCCTACACGACGCTCTTCCGATCTNNNNCGTATGCAAGCCCTGGACCAGGCA |
| P5_IgD_A8 | ACACTCTTTCCCTACACGACGCTCTTCCGATCTNNNNCGTATGCACCCTGATATGATGGGGAACA |
| P5_IgM_A8 | ACACTCTTTCCCTACACGACGCTCTTCCGATCTNNNNCGTATGCAGGGAATTCTCACAGGAGACG |
| P5_IgE_A8 | ACACTCTTTCCCTACACGACGCTCTTCCGATCTNNNNCGTATGCAGAAGACGGATGGGCTCTGT |
| P5_IgG_long_A9 | ACACTCTTTCCCTACACGACGCTCTTCCGATCTNNNNCGAGCAGACAGGCAGCCCAGGGC |
| P5_IgA_long_A9 | ACACTCTTTCCCTACACGACGCTCTTCCGATCTNNNNCGAGCAGAAGCCCTGGACCAGGCA |

| | |
|---|---|
| P5_IgD_A9 | ACACTCTTTCCCTACACGACGCTCTTCCGATCTNNNNCGAGCA GACCCTGATATGATGGGGAACA |
| P5_IgM_A9 | ACACTCTTTCCCTACACGACGCTCTTCCGATCTNNNNCGAGCA GAGGGAATTCTCACAGGAGACG |
| P5_IgE_A9 | ACACTCTTTCCCTACACGACGCTCTTCCGATCTNNNNCGAGCA GAGAAGACGGATGGGCTCTGT |
| P5_IgG_long_A10 | ACACTCTTTCCCTACACGACGCTCTTCCGATCTNNNNGATCGA GCCAGGCAGCCCAGGGC |
| P5_IgA_long_A10 | ACACTCTTTCCCTACACGACGCTCTTCCGATCTNNNNGATCGA GCAGCCCTGGACCAGGCA |
| P5_IgD_A10 | ACACTCTTTCCCTACACGACGCTCTTCCGATCTNNNNGATCGA GCCCCTGATATGATGGGGAACA |
| P5_IgM_A10 | ACACTCTTTCCCTACACGACGCTCTTCCGATCTNNNNGATCGA GCGGGAATTCTCACAGGAGACG |
| P5_IgE_A10 | ACACTCTTTCCCTACACGACGCTCTTCCGATCTNNNNGATCGA GCGAAGACGGATGGGCTCTGT |
| P5_IgG_long_A11 | ACACTCTTTCCCTACACGACGCTCTTCCGATCTNNNNGCATGC ACCAGGCAGCCCAGGGC |
| P5_IgA_long_A11 | ACACTCTTTCCCTACACGACGCTCTTCCGATCTNNNNGCATGC ACAGCCCTGGACCAGGCA |
| P5_IgD_A11 | ACACTCTTTCCCTACACGACGCTCTTCCGATCTNNNNGCATGC ACCCCTGATATGATGGGGAACA |
| P5_IgM_A11 | ACACTCTTTCCCTACACGACGCTCTTCCGATCTNNNNGCATGC ACGGGAATTCTCACAGGAGACG |
| P5_IgE_A11 | ACACTCTTTCCCTACACGACGCTCTTCCGATCTNNNNGCATGC ACGAAGACGGATGGGCTCTGT |
| P5_IgG_long_A12 | ACACTCTTTCCCTACACGACGCTCTTCCGATCTNNNNTACGAT GCCAGGCAGCCCAGGGC |
| P5_IgA_long_A12 | ACACTCTTTCCCTACACGACGCTCTTCCGATCTNNNNTACGAT GCAGCCCTGGACCAGGCA |
| P5_IgD_A12 | ACACTCTTTCCCTACACGACGCTCTTCCGATCTNNNNTACGAT GCCCCTGATATGATGGGGAACA |
| P5_IgM_A12 | ACACTCTTTCCCTACACGACGCTCTTCCGATCTNNNNTACGAT GCGGGAATTCTCACAGGAGACG |
| P5_IgE_A12 | ACACTCTTTCCCTACACGACGCTCTTCCGATCTNNNNTACGAT GCGAAGACGGATGGGCTCTGT |

### References


1. K. M. Roskin *et al.*, IgH sequences in common variable immune deficiency reveal altered B cell development and selection. *Sci Transl Med* **7**, 302ra135 (2015).
2. J. Ye, N. Ma, T. L. Madden, J. M. Ostell, IgBLAST: an immunoglobulin variable domain sequence analysis tool. *Nucleic Acids Res* **41**, W34-40 (2013).

3. M. P. Lefranc, IMGT, the international ImMunoGeneTics database. *Nucleic Acids Res* **31**, 307-310 (2003).
4. C. Wang *et al.*, B-cell repertoire responses to varicella-zoster vaccination in human identical twins. *Proc Natl Acad Sci U S A* **112**, 500-505 (2015).
5. M. P. Lefranc *et al.*, IMGT(R), the international ImMunoGeneTics information system(R) 25 years on. *Nucleic Acids Res* **43**, D413-422 (2015).
6. S. C. A. Nielsen *et al.*, Shaping of infant B cell receptor repertoires by environmental factors and infectious disease. *Sci Transl Med* **11**, (2019).
7. A. H. Lucas *et al.*, Molecular ontogeny of the human antibody repertoire to the Haemophilus influenzae type B polysaccharide: expression of canonical variable regions and their variants in vaccinated infants. *Clin Immunol* **108**, 119-127 (2003).
8. J. Truck *et al.*, Identification of antigen-specific B cell receptor sequences using public repertoire analysis. *J Immunol* **194**, 252-261 (2015).
9. E. E. Adderson *et al.*, Restricted immunoglobulin VH usage and VDJ combinations in the human response to Haemophilus influenzae type b capsular polysaccharide. Nucleotide sequences of monospecific anti-Haemophilus antibodies and polyspecific antibodies cross-reacting with self antigens. *J Clin Invest* **91**, 2734-2743 (1993).
10. A. H. Lucas, J. W. Larrick, D. C. Reason, Variable region sequences of a protective human monoclonal antibody specific for the Haemophilus influenzae type b capsular polysaccharide. *Infect Immun* **62**, 3873-3880 (1994).
11. J. D. Berry, D. J. Boese, D. K. Law, W. D. Zollinger, R. S. Tsang, Molecular analysis of monoclonal antibodies to group variant capsular polysaccharide of Neisseria meningitidis: recurrent heavy chains and alternative light chain partners. *Mol Immunol* **42**, 335-344 (2005).
12. S. L. Smithson, N. Srivastava, W. A. Hutchins, M. A. Westerink, Molecular analysis of the heavy chain of antibodies that recognize the capsular polysaccharide of Neisseria meningitidis in hu-PBMC reconstituted SCID mice and in the immunized human donor. *Mol Immunol* **36**, 113-124 (1999).
13. W. A. Hutchins, A. R. Adkins, T. Kieber-Emmons, M. A. Westerink, Molecular characterization of a monoclonal antibody produced in response to a group C meningococcal polysaccharide peptide mimic. *Mol Immunol* **33**, 503-510 (1996).
14. Z. Chen *et al.*, Human monoclonal antibodies isolated from a primary pneumococcal conjugate Vaccinee demonstrates the expansion of an antigen-driven Hypermutated memory B cell response. *BMC Infect Dis* **18**, 613 (2018).
15. K. Kolibab, S. L. Smithson, B. Rabquer, S. Khuder, M. A. Westerink, Immune response to pneumococcal polysaccharides 4 and 14 in elderly and young adults: analysis of the variable heavy chain repertoire. *Infect Immun* **73**, 7465-7476 (2005).
16. K. Smith *et al.*, Fully human monoclonal antibodies from antibody secreting cells after vaccination with Pneumovax(R)23 are serotype specific and facilitate opsonophagocytosis. *Immunobiology* **218**, 745-754 (2013).
17. A. S. Adler *et al.*, Rare, high-affinity anti-pathogen antibodies from human repertoires, discovered using microfluidics and molecular genomics. *MAbs* **9**, 1282-1296 (2017).
18. B. J. DeKosky *et al.*, High-throughput sequencing of the paired human immunoglobulin heavy and light chain repertoire. *Nature biotechnology* **31**, 166-169 (2013).

19. J. de Kruif *et al.*, Human immunoglobulin repertoires against tetanus toxoid contain a large and diverse fraction of high-affinity promiscuous V(H) genes. *J Mol Biol* **387**, 548-558 (2009).
20. D. Frolich *et al.*, Secondary immunization generates clonally related antigen-specific plasma cells and memory B cells. *J Immunol* **185**, 3103-3110 (2010).
21. T. R. Poulsen, A. Jensen, J. S. Haurum, P. S. Andersen, Limits for antibody affinity maturation and repertoire diversification in hypervaccinated humans. *J Immunol* **187**, 4229-4235 (2011).
22. M. S. Gilman *et al.*, Rapid profiling of RSV antibody repertoires from the memory B cells of naturally infected adult donors. *Sci Immunol* **1**, (2016).
23. J. V. Williams, J. H. Weitkamp, D. L. Blum, B. J. LaFleur, J. E. Crowe, Jr., The human neonatal B cell response to respiratory syncytial virus uses a biased antibody variable gene repertoire that lacks somatic mutations. *Mol Immunol* **47**, 407-414 (2009).
24. A. Tang *et al.*, A potent broadly neutralizing human RSV antibody targets conserved site IV of the fusion glycoprotein. *Nat Commun* **10**, 4153 (2019).
25. B. Cortjens *et al.*, Broadly Reactive Anti-Respiratory Syncytial Virus G Antibodies from Exposed Individuals Effectively Inhibit Infection of Primary Airway Epithelial Cells. *J Virol* **91**, (2017).
26. E. Goodwin *et al.*, Infants Infected with Respiratory Syncytial Virus Generate Potent Neutralizing Antibodies that Lack Somatic Hypermutation. *Immunity* **48**, 339-349 e335 (2018).
27. J. C. Krause *et al.*, Epitope-specific human influenza antibody repertoires diversify by B cell intraclonal sequence divergence and interclonal convergence. *J Immunol* **187**, 3704-3711 (2011).
28. M. A. Moody *et al.*, H3N2 influenza infection elicits more cross-reactive and less clonally expanded anti-hemagglutinin antibodies than influenza vaccination. *PloS one* **6**, e25797 (2011).
29. J. Wrammert *et al.*, Broadly cross-reactive antibodies dominate the human B cell response against 2009 pandemic H1N1 influenza virus infection. *J Exp Med* **208**, 181-193 (2011).
30. A. H. Ellebedy *et al.*, Defining antigen-specific plasmablast and memory B cell subsets in human blood after viral infection or vaccination. *Nat Immunol* **17**, 1226-1234 (2016).
31. D. Corti *et al.*, Tackling influenza with broadly neutralizing antibodies. *Curr Opin Virol* **24**, 60-69 (2017).
32. K. E. Neu *et al.*, Spec-seq unveils transcriptional subpopulations of antibody-secreting cells following influenza vaccination. *J Clin Invest* **129**, 93-105 (2019).
33. Y. Liu *et al.*, Cross-lineage protection by human antibodies binding the influenza B hemagglutinin. *Nat Commun* **10**, 324 (2019).
34. D. Hirano *et al.*, Three Types of Broadly Reacting Antibodies against Influenza B Viruses Induced by Vaccination with Seasonal Influenza Viruses. *J Immunol Res* **2018**, 7251793 (2018).
35. A. Yasuhara *et al.*, Diversity of antigenic mutants of influenza A(H1N1)pdm09 virus escaped from human monoclonal antibodies. *Scientific reports* **7**, 17735 (2017).
36. C. Shen *et al.*, A multimechanistic antibody targeting the receptor binding site potently cross-protects against influenza B viruses. *Sci Transl Med* **9**, (2017).

37. M. G. Joyce *et al.*, Vaccine-Induced Antibodies that Neutralize Group 1 and Group 2 Influenza A Viruses. *Cell* **166**, 609-623 (2016).
38. Y. Fu *et al.*, A broadly neutralizing anti-influenza antibody reveals ongoing capacity of haemagglutinin-specific memory B cells to evolve. *Nat Commun* **7**, 12780 (2016).
39. G. M. Li *et al.*, Pandemic H1N1 influenza vaccine induces a recall response in humans that favors broadly cross-reactive memory B cells. *Proceedings of the National Academy of Sciences of the United States of America* **109**, 9047-9052 (2012).
40. A. Watanabe *et al.*, Antibodies to a Conserved Influenza Head Interface Epitope Protect by an IgG Subtype-Dependent Mechanism. *Cell* **177**, 1124-1135 e1116 (2019).
41. K. R. McCarthy *et al.*, Memory B Cells that Cross-React with Group 1 and Group 2 Influenza A Viruses Are Abundant in Adult Human Repertoires. *Immunity* **48**, 174-184 e179 (2018).
42. S. F. Andrews *et al.*, Preferential induction of cross-group influenza A hemagglutinin stem-specific memory B cells after H7N9 immunization in humans. *Sci Immunol* **2**, (2017).
43. P. J. M. Brouwer *et al.*, Potent neutralizing antibodies from COVID-19 patients define multiple targets of vulnerability. *Science*, (2020).
44. A. Z. Wec *et al.*, Broad neutralization of SARS-related viruses by human monoclonal antibodies. *Science*, (2020).
45. T. F. Rogers *et al.*, Isolation of potent SARS-CoV-2 neutralizing antibodies and protection from disease in a small animal model. *Science*, (2020).
46. S. J. Zost *et al.*, Rapid isolation and profiling of a diverse panel of human monoclonal antibodies targeting the SARS-CoV-2 spike protein. *Nat Med*, (2020).
47. R. Shi *et al.*, A human neutralizing antibody targets the receptor-binding site of SARS-CoV-2. *Nature*, (2020).
48. E. Seydoux *et al.*, Characterization of neutralizing antibodies from a SARS-CoV-2 infected individual. *bioRxiv*, (2020).
49. D. F. Robbiani *et al.*, Convergent Antibody Responses to SARS-CoV-2 Infection in Convalescent Individuals. *bioRxiv*, (2020).
50. D. Pinto *et al.*, Structural and functional analysis of a potent sarbecovirus neutralizing antibody. *bioRxiv*, (2020).
51. C. Kreer *et al.*, Longitudinal Isolation of Potent Near-Germline SARS-CoV-2-Neutralizing Antibodies from COVID-19 Patients. *Cell*, (2020).
52. C. W. Davis *et al.*, Longitudinal Analysis of the Human B Cell Response to Ebola Virus Infection. *Cell* **177**, 1566-1582 e1517 (2019).
53. P. Virtanen *et al.*, SciPy 1.0: fundamental algorithms for scientific computing in Python. *Nature methods* **17**, 261-272 (2020).
54. L. Fu, B. Niu, Z. Zhu, S. Wu, W. Li, CD-HIT: accelerated for clustering the next-generation sequencing data. *Bioinformatics* **28**, 3150-3152 (2012).
55. W. McKinney, paper presented at the Proceedings of the 9th Python in Science Conference, 2010.
56. R Development Core Team. (R Foundation for Statistical Computing, Vienna, Austria, 2010).
